# Coiled Coil Crosslinked Alginate Hydrogels Dampen Macrophage-Driven Inflammation

**DOI:** 10.1101/2021.11.01.466776

**Authors:** Zain Clapacs, Conor O’Neill, Paresh Shrimali, Giriraj Lokhande, Megan Files, Darren D. Kim, Akhilesh Gaharwar, Jai S. Rudra

## Abstract

Alginate hydrogels are widely used for tissue engineering and regenerative medicine due to their excellent biocompatibility. A facile and commonly used strategy to crosslink alginate is the addition of Ca^2+^ that leads to hydrogelation. However, extracellular Ca^2+^ is a secondary messenger in activating inflammasome pathways following physical injury or pathogenic insult leading to persistent inflammation and scaffold rejection. Here we present graft copolymers of charge complementary heterodimeric coiled coil (CC) peptides and alginate that undergo supramolecular self-assembly to form Ca^2+^ free alginate hydrogels. The formation of heterodimeric CCs was confirmed using circular dichroism spectroscopy and scanning electron microscopy revealed a significant difference in pore size between Ca^2+^ and CC crosslinked gels. The resulting hydrogels were self-supporting and display shear-thinning and shear-recovery properties. In response to lipopolysaccharide (LPS) stimulation, peritoneal macrophages and bone marrow derived dendritic cells cultured in the CC crosslinked gels exhibited a 10-fold reduction in secretion of the proinflammatory cytokine IL-1β compared to Ca^2+^ crosslinked gels. A similar respose was also observed *in vivo* upon peritoneal delivery of Ca^2+^ or CC crosslinked gels. Analysis of peritoneal lavage showed that macrophages in mice injected with Ca^2+^ crosslinked gels display a more inflammatory phenotype compared to macrophages from mice injected with CC crosslinked gels. These results suggest that CC peptides by virtue of their tunable sequence-structure-function relationship and mild gelation conditions are promising alternative crosslinkers for alginate and other biopolymer scaffolds used in tissue engineering.

## Introduction

Promoting a proper immune response to scaffold materials is of paramount importance for tissue engineers to avoid rejection and allow native-like tissue regrowth *in vivo*.^1–6^ In particular, macrophages (MΦs) and their immunological response to scaffold materials are of interest due to their influence on wound healing and tissue remodeling processes.^2,7,8^ Although MΦ-driven inflammation is necessary and beneficial in the early stages of wound healing, failure of MΦs to subsequently transition from a pro-inflammatory to pro-proliferative phenotype can lead to chronic inflammation at the wound site.^8,9^ Previous studies have shown that MΦ driven production of the pro-inflammatory cytokine interleukin-1β (IL-1β) is a major mechanism contributing to such failure to transition, particularly in non-healing wounds.^8,10^ For this reason, it is important that the materials and excipients used in scaffolds aimed at tissue regeneration avoid MΦ-driven production of IL-1β, which could lead to persistent inflammation and scaffold rejection.

Hydrogels based on diverse natural and synthetic polymers have been used as scaffolds for tissue engineering.^11–15^ Alginate hydrogels in particular are notable for their biocompatibility with somatic and multipotent cells, robust mechanical properties, mild conditions of gelation, ease of modification, and structural similarities to extracellular matrix.^11,16–19^ Although alginate is easy to crosslink non-covalently with the addition of Ca^2+^ ions, Ca^2+^-alginate gels have been shown *in vivo* to leach Ca^2+^ into the physiological environment through cation exchange.^16,20^ In particular this presents difficulties because extracellular Ca^2+^ is known to stimulate activation of the NLRP3 inflammasome pathway via stimulation of G protein-coupled calcium sensing receptors.^21^ This pro-inflammatory effect has been observed both *in vitro* and *in vivo* leading to elevated expression of IL-1β.^22^ This has led to an interest in Ca^2+^-free methods of crosslinking alginate, however covalently crosslinked hydrogels have presented some difficulties related to injectability, as well as reduced ease of use compared to conventional Ca^2+^ crosslinking^17^ while noncovalent methods involving harsh solvents, metal ions, cryogelation and phase separation have found limited biological application.^23–26^ These hurdles present a need for a biocompatible and noncovalent method for crosslinking alginate while avoiding the inflammatory response characteristic of Ca^2+^ crosslinked gels.

Coiled coil peptides (CCs), which form a stable supramolecular structure of supercoiled α-helices have been shown to be viable crosslinkers for synthetic polymers, both through metal-ion induced assembly in-situ and through the mixing of complimentary graft copolymers^27–30^ and recent work has demonstrated the utility of CCs to crosslink hyaluronan.^31^ Previous investigations have largely focused on CC crosslinked gels at a microscale in the context of drug delivery, with relatively little investigation of their potential to form bulk hydrogels for cell encapsulation.^29,31^ Notably, the immunogical effects of CC end-linked supramolecular hydrogels are minimal and *in vivo* such hydrogels do not induce substantial cytokine responses.^32–34^ In this work we demonstrate the use of CCs as a mechanism to noncovalently crosslink alginate hydrogels in the absence of Ca^2+^. We demonstrate that hydrogels formed by alginate-CC graft copolymers (**Figure 1**) can support encapsulation of primary macrophages (MΦs) and dendritic cells (DCs) with minimal IL-1β production compared to Ca^2+^ crosslinked gels. We also demonstrate that peritoneal MΦs isolated from mice injected with CC crosslinked gels display a milder inflammatory phenotype than MΦs isolated from mice injected with Ca^2+^ crosslinked alginate.

**Scheme 1.**
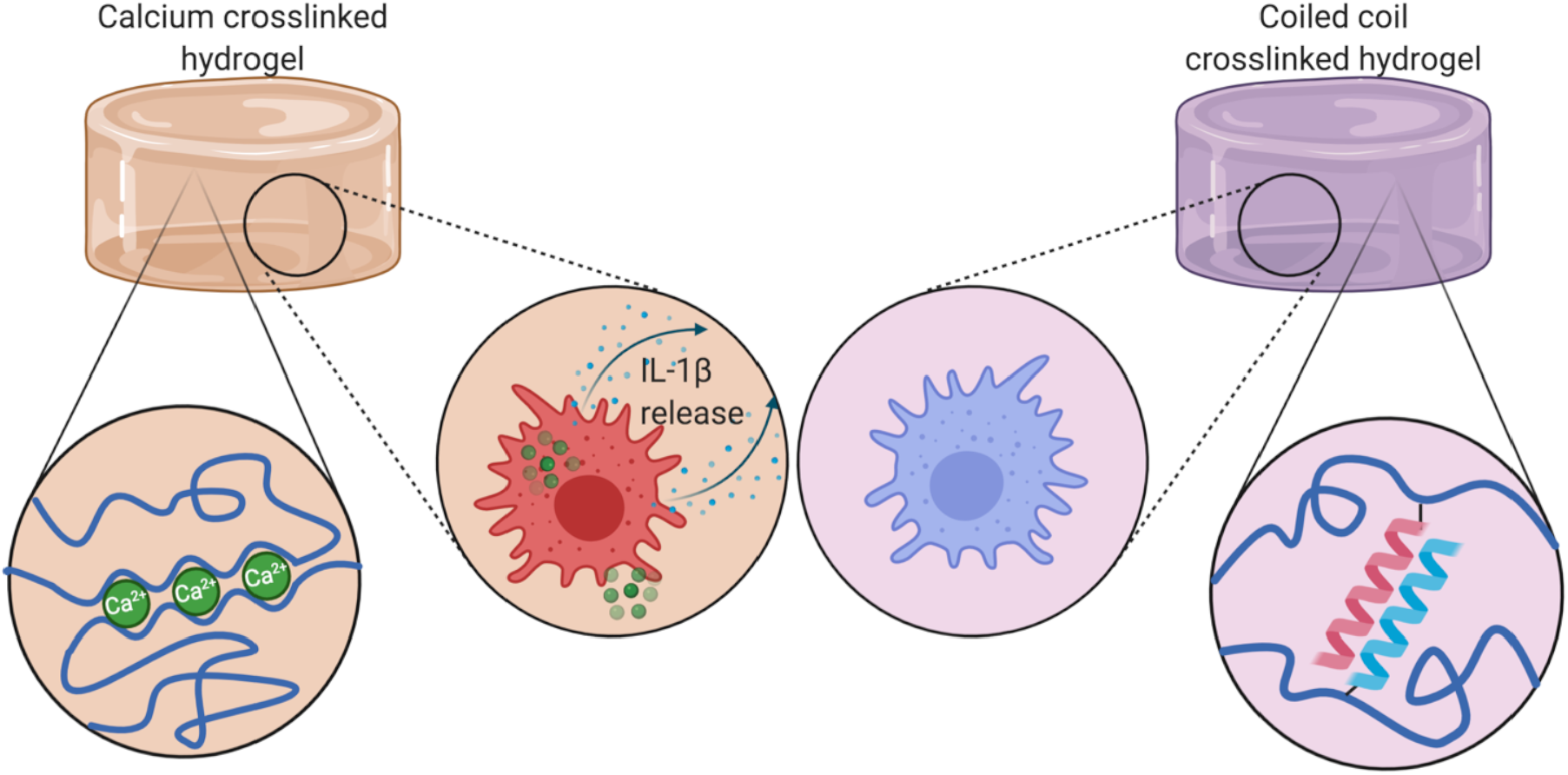
Schematic of Ca^2+^ and CC crosslinking in alginate hydrogels. Ca^2+^ crosslinked hydrogels leach Ca^2+^ ions into the environment which trigger IL-1β release.

**Figure 1:**
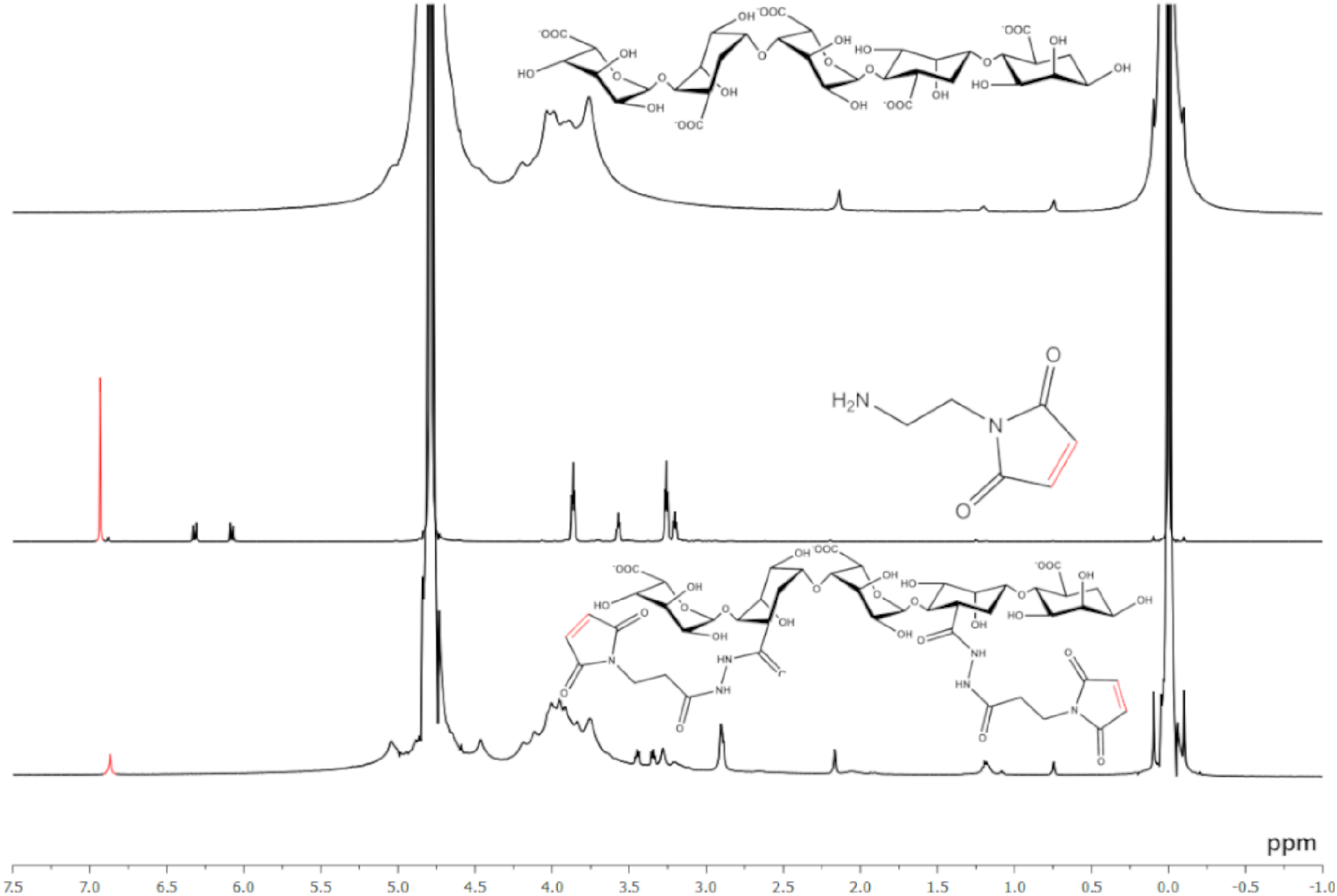
^1^H NMR spectra of alginate, maleimidoethylamine, and BMPH conjugated alginate in D_2_O, 600 MHz. Peaks associated with maleimide moieties highlighted in red. Spectrum annotation in (**Figure S6-S8**).

## Experimental Section/Methods

### Materials

Fmoc-amino acids and oxyma were purchased from Chem-Impex. Amino-acid preloaded Wang resins for peptide synthesis were purchased from Novabiochem. Diisopropylcarbodiimide, triisopropylsilane and ethanedithiol were purchased from Millipore. Trifluoroacetic acid and N-β-maleimidopropionic acid hydrazide (BMPH) were purchased from Thermo Fisher. Piperidine was purchased from Alfa Asear. N,N-dimethylformamide was purchased from Arcos Organics. Sodium alginate, alginate lyase, RPMI-1640 media in powder form and 2-(N-morpholino)ethanesulfonic acid were purchased from Sigma Aldrich. 1-ethyl-3-(3-dimethylaminopropyl) carbodiimide was Purchased from G-Biosciences.

### Graft copolymer synthesis

E3 and K3 peptides (sequences **Table S1**) were synthesized using solid phase peptide synthesis (**Scheme S1**) assisted by a Liberty Blue automated peptide synthesizer (CEM) and purified using high performance liquid chromatograpy (HPLC) (**Figure S1-S3**). The identity of the purified peptides was confirmed using MALDI-TOF mass spectrometry (**Figure S4-S5**). The peptides were grafted to alginate by first ligating BMPH to alginate using carbodiimide mediated amine-carboxyl ligation. The free maleimide moiety of BMPH was used couple the peptide via N-terminal cysteine residues on E3 or K3. Detailed methods are provided in SI.

### Material characterization

Ligation of BMPH to alginate was confirmed using ^1^H NMR and maleimide-thiol reaction efficiency was quantified using FAM-thiol as a surrogate. E3/K3 conjugation was confirmed and quantified using BCA analysis. Circular dichroism (CD) spectroscopy of the free peptides and peptide-alginate conjugates was used to confirm coiled coil formation. Microporous structure was investigated using scanning electron microscopy (SEM) after flash freezing samples by submerging in liquid nitrogen and lyophilizing. The shear response, strain recovery and temperature sensitivity was assessed using parallel plate rheology and their ability to support 3D cell cultures was confirmed using mouse peritoneal MΦs. Detailed methods are provided in SI.

### In vitro cellular response

Peritoneal MΦs were isolated from mice and encapsulated in either CC or Ca^2+^ crosslinked gels and stimulated for 24 hours with 500 ng/mL LPS. To assess IL-1β secretion supernatant was collected and analyzed using enzyme linked immunosorbent assay (ELISA). Cell phenotype was assessed by degrading the gels using alginate lyase and staining cells for the surface markers F4/80, MHC-II, CD40, CD80 and CD86 and flow cytometry analysis. Detailed methods are provided in SI.

### In vivo immunogenicity

All animal work was approved by the Institutional Animal Care and Use Committee at Washington University in St. Louis. Animals were provided food and water ad libitium and kept in a climate controlled facility with a 12 hour light/dark cycle. 6-week old C57BL6/J mice were divided into 4 groups of 3 and each group was intraperitoneally administered a 150 μL injection of either a 0.5% alginate solution or a 0.5% alginate hydrogel crosslinked either with CCs, CaCl_2_ or CaSO_4_. After 24 hours animals were euthanized and peritoneal cells were collected by peritoneal lavage. Cells were pooled within treatment groups and stained for CD11b, F4/80, Ly6-G, MHC-II, and a nuclear stain for dead cell exclusion. Cells were then analyzed using flow cytometry. Detailed methods are provided in SI.

## Results & Discussion

### Graft Copolymer Synthesis

Alginate hydrogels present many benefits for tissue engineering due to their biocompatibility, immunological inertness, and versatility for chemical modification with bioactive moieties. However due to its polyanionic nature, alginate is susceptible to charge-charge aggregation with polycations or positively charged peptides and presents a challenge for the use of charge-complementary CCs.^35–37^ To overcome this, we employed a similar strategy to that described by Ding et al for hyaluronic acid.^31^ We used a modified version of the previously described E3/K3 system (**Sequences in Table S1**). Notably, because the coiled coil forming region in the E3/K3 system is only 3 heptads long, the net charge on the cationic peptide (K3) is only +3 which improves the resilience of the alginate-K3 graft copolymer to aggregation.^38^ Further, the interaction of E3/K3 can be specifically detected using circular dichroism (CD) spectroscopy because the peptides only exhibit a helical trace when supercoiled.^31,39^ To construct the alginate-peptide graft copolymer, alginate was functionalized with the bifunctional crosslinker BMPH which presents a thiol-reactive maleimide group and a free hydrazide, which was then linked to a free cysteine residue presented on the N-terminus of either the E3 or K3 peptide (**Scheme 2**).

**Scheme 2:**
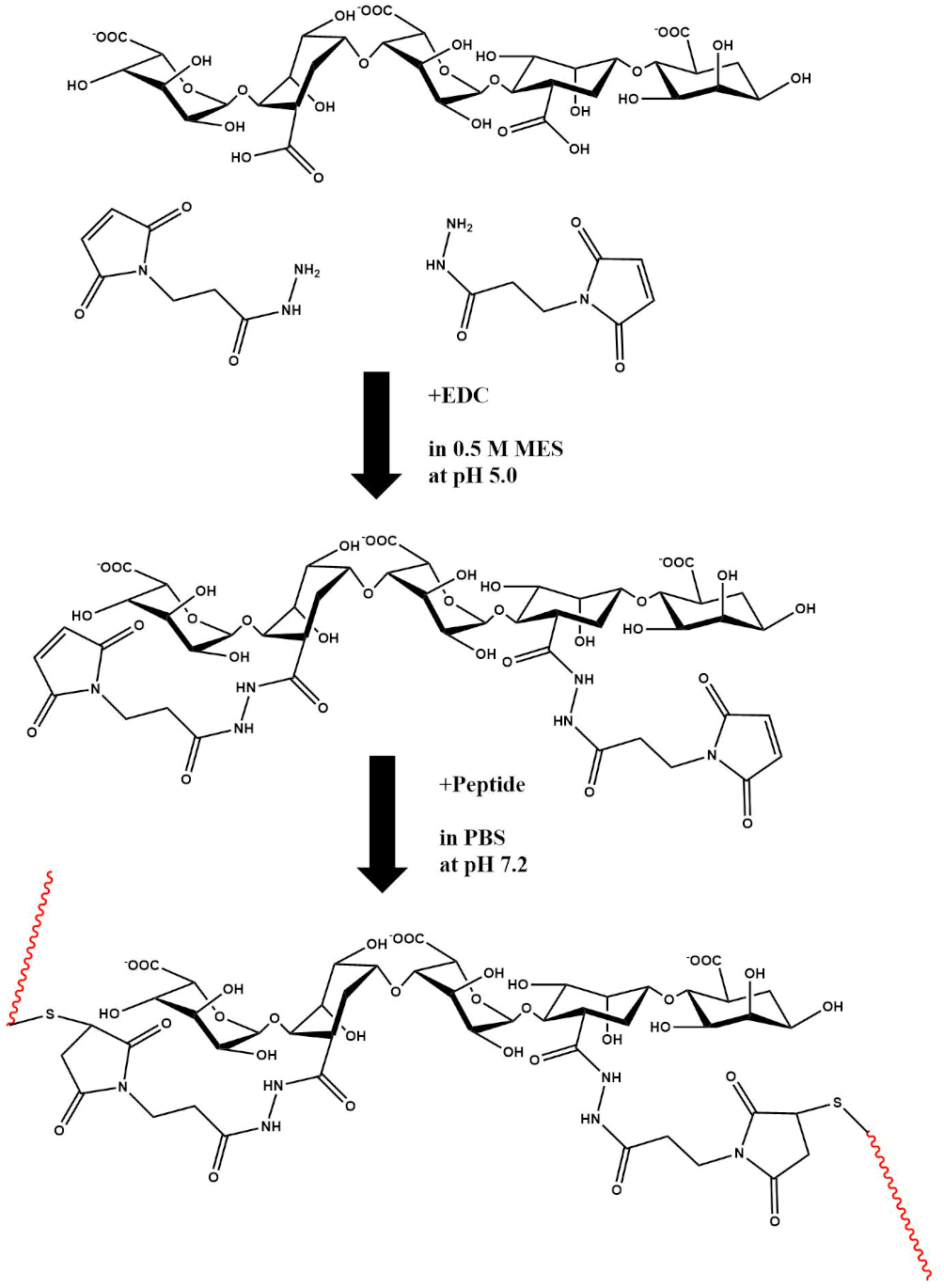
Synthesis of peptide-alginate graft co-polymer. Alginate was functionalized with either E3 or K3 peptide via a 2 step process. First, BMPH was ligated to alginate using carbodiimide mediated amine-carboxyl ligation, then the peptide was bound to the maleimide moiety of BMPH at a cysteine residue through maleimide-thiol thioether formation.

We performed ^1^H NMR spectroscopy on the product to confirm the presence of maleimide groups to ensure reaction completion (**Figure 1**). NMR spectra were obtained for alginate, maleimidoethylamine as a model maleimide presenting compound and 2% BMPH substitution alginate. The presence of maleimide after carbodiimide ligation was confirmed by the presence of a singlet peak at δ=6.87 ppm in the spectrum of the BMPH-alginate conjugate, a peak which has been previously characterized as corresponding to maleimides and similar to the most downfield peak observed in the maleimidoethylamine trace.^40^

Three alginate-BMPH conjugates were made at theoretical molar substitution rates of 1%, 2% and 5% BMPH/uronic acid to investigate the degree to which alginate could be functionalized with K3 before inducing charge aggregation. Alginate-BMPH at 1%, 2% and 5% BMPH substitution was then reacted with a 2x molar excess of K3 overnight (**Scheme 2**) and unreacted maleimide groups were quenched with an excess of free cysteine. The 5% theoretical substitution alginate was observed to have formed insoluble aggregates while the other two substitutions did not, for that reason all further work was conducted with the 2% theoretical substitution alginate as it was the greatest substitution of K3 peptide to alginate that was not intractable. The reaction efficiency of peptide substitution was quantified by reacting alginate-BMPH (2% theoretical substitution) with FAM-thiol at the same conditions as K3 peptide. Data indicated a final molar substitution of 1.54% corresponding to 77% total synthesis efficiency (**Figure 2**). These results were confirmed via bichinchoninic acid assay. E3 peptide was conjugated to alginate-BMPH (2%) and all subsequent work was conducted using the graft copolymers.

**Figure 2.**
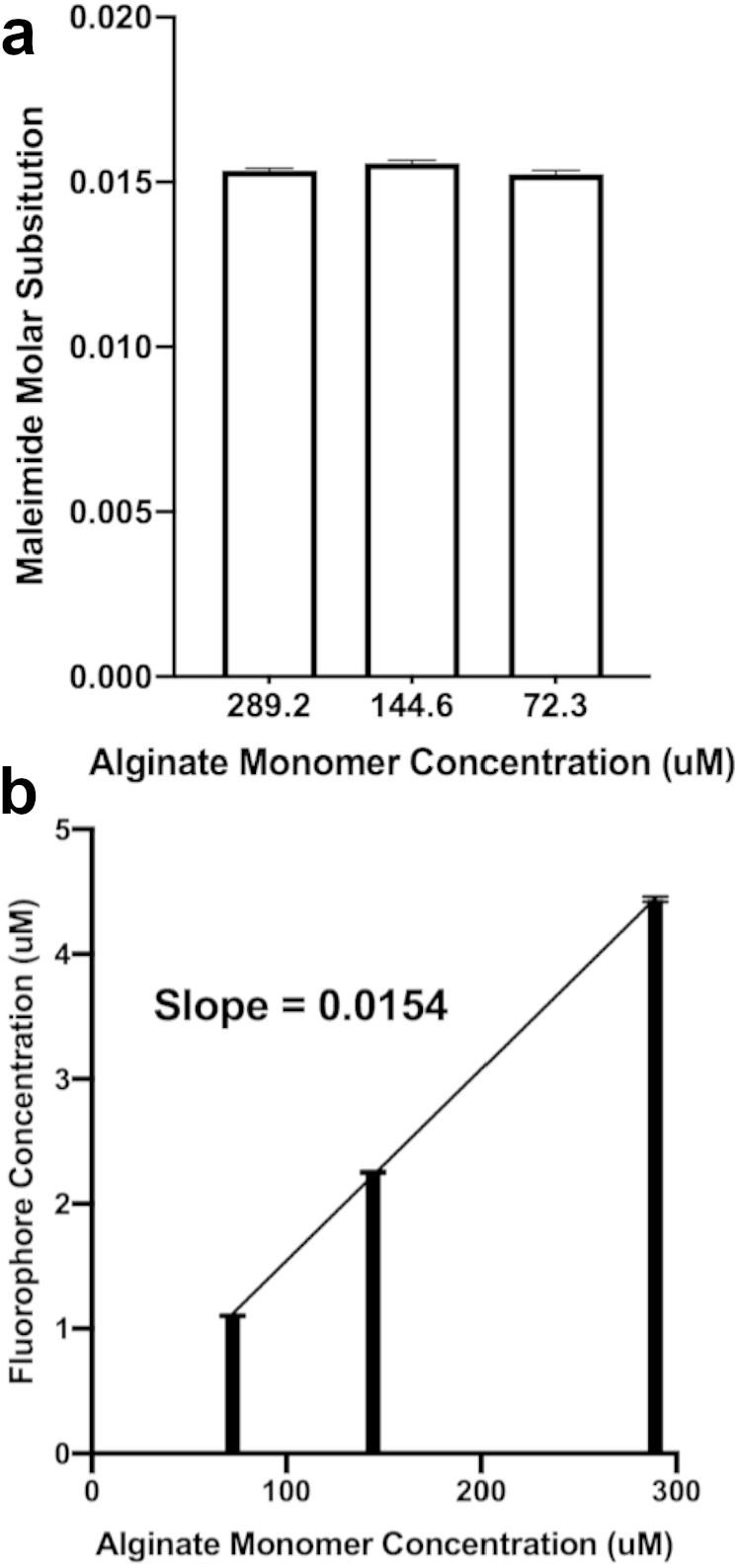
Quantification of BMPH-alginate functionalization. Final conjugation capacity was assessed by conjugation of FAM-thiol to BMPH-alginate under identical conditions to those used for peptide conjugation to BMPH-alginate, quantitative fluorescence analysis demonstrated final molar substitution approximated at (a) individual concentrations and (b) between concentrations of approximately 1.54%.

### Crosslink Characterization

When dissolved separately in PBS at 0.5% w/v both alginate-E3 and alginate-K3 were observed to flow as viscous fluids, and when equal volumes of the solutions were mixed by pipetting, they formed a self-supporting gel (**Figure S9**). As the coiled coil forming domains of the E3/K3 heterodimer only adopt an α-helical conformation when stabilized, CD is an effective method to investigate if the observed gelation is due to supramolecular self-assembly or simple charge-aggregation (alginate and PBS blank spectra **Figure S10**).^39^ The ratio of the peaks at 222 and 208 nm can be used as a measurement of helicity, as a high ratio [θ]_220_/[θ]_208_ (referred to as the helicity ratio) indicates a more coiled coil-like conformation ^41^ and generally helicity ratios >0.9 indicate potential coiled coil formation.^41–43^ This ratio can also be used to confirm that a mixture of peptides structurally interacts in solution. Because the spectrum of a mixture of non-interacting molecules is a linear combination of the components’ spectra, a mixture with a spectrum different than the expected indicates that the components structurally interact.^44^ Therefore, a mixture with a helicity ratio substantially different from the number average helicity ratio of its components contains components that change conformation into a more or less coiled coil-like structure when mixed. The spectrum of free K3 displayed characteristic features of a random coil while free E3 and an equimolar mixture of E3 and K3 (E3/K3) had α-helical spectra (**Figure 3a**). The α-helical spectrum of E3 likely corresponds to the formation of a previously observed unstable homotrimer (**Figure 3e**).^39^ The free E3/K3 mixture also displayed a helical trace, in this case likely corresponding to formation of the intended heterodimeric coiled coil. This is further illustrated by the helicity ratios of free E3, free K3 and free E3/K3 mixture (**Figure 3c**). If the helicity observed in the mixture were driven primarily by the presence of the E3 homotrimer then the observed helicity of the mixture would be approximately equal to the average of E3 and K3’s helicity ratios when dissolved independently, however the observed helicity ratio of free E3/K3 mixture is much larger indicating interaction between E3 and K3 leading both peptides to adopt a helical conformation, as would be expected in formation of the E3/K3 heterodimer (**Figure 4f**). When conjugated to alginate, the observed spectra of the E3 and K3 peptides alone changed markedly. Alginate-E3 displayed a more random coil-like trace while alginate-K3 and the equimolar mixture of alginate-E3 and alginate-K3 (alginate-E3/K3) displayed helical traces. (**Figure 3b**). The loss of helicity in E3 when conjugated to alginate may be due to steric disruption of the already unstable homotrimer. On the contrary, improved helicity of alginate-K3 may be due to charge stabilization of the cationic peptide by alginate (**Figure 3d**). Similarly to the case of free peptides, the alginate-E3/K3 mixture had a helicity ratio substantially greater than the average of the components’ ratios, indicating that alginate-E3 and alginate-K3 similarly interact in solution to form a heterodimeric coiled coil.

**Figure 3.**
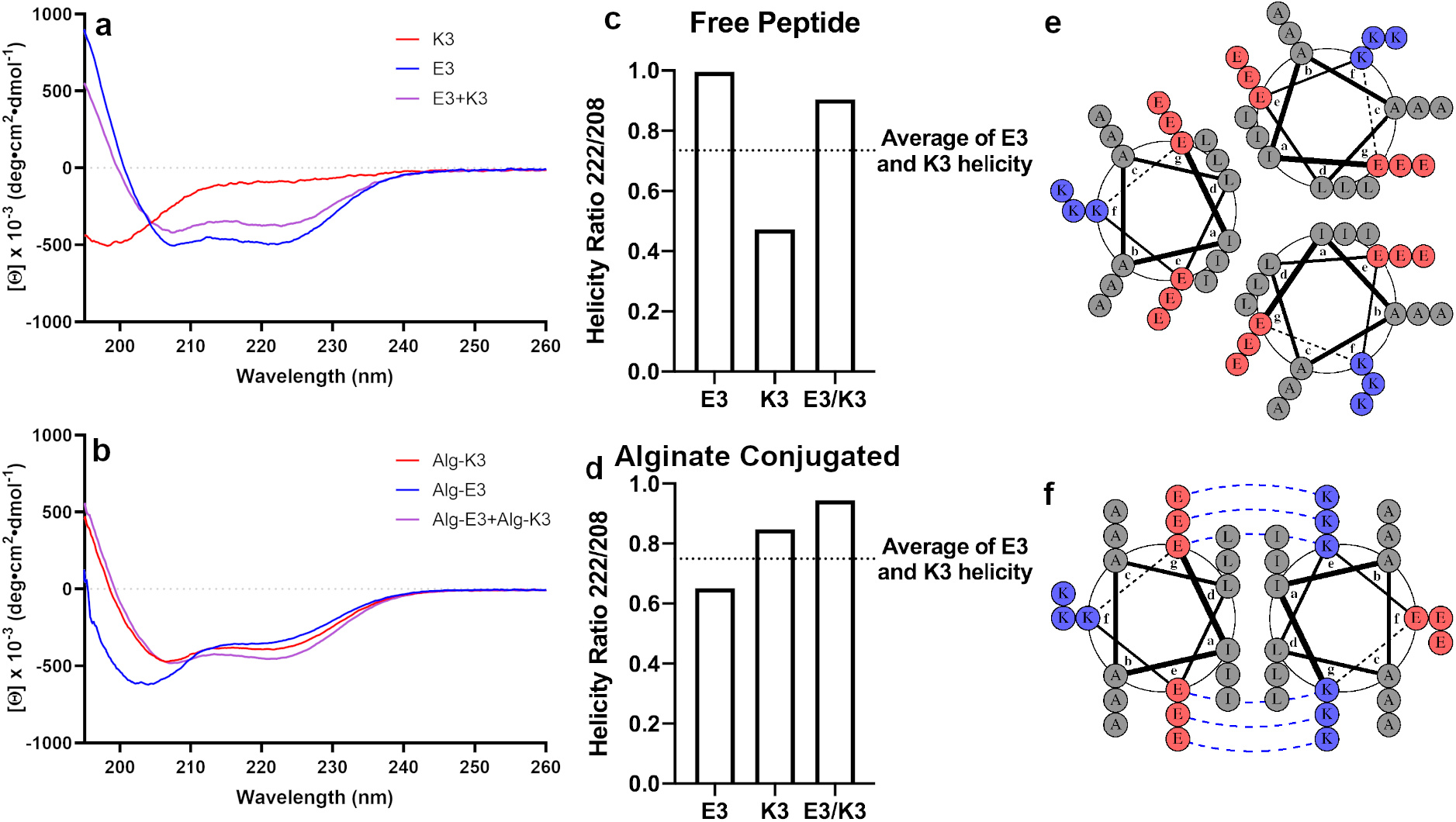
Circular dichroism analysis of free and alginate-conjugated E3/K3 peptides. (a,b) Circular dichroism spectroscopy of free (a) and alginate conjugated (b) peptides indicates that the blended solution does form helical structures. Notably, in free solution E3 also displays a helical trace which may correspond to E3 homotrimers (**Figure 3e**). While in alginate conjugated solution K3 displays a helical trace which may correspond to charge based stabilization between the positively charged peptide and negatively charged alginate. (c,d) Both free (c) and alginate conjugated (d) solutions of E3/K3 blends display a much greater helicity than would be expected if the peptides did not interact indicating formation the expected coiled-coil dimer. (e) The helical trace observed in free E3 (**Figure 3a**) may correspond to a homotrimeric coiled coil previously described by Apostolovic & Klok.^39^ (f) The helical trace in blended solutions (**Figure 3a**,**b**) indicates the formation of the E3/K3 heterodimer.

**Figure 4.**
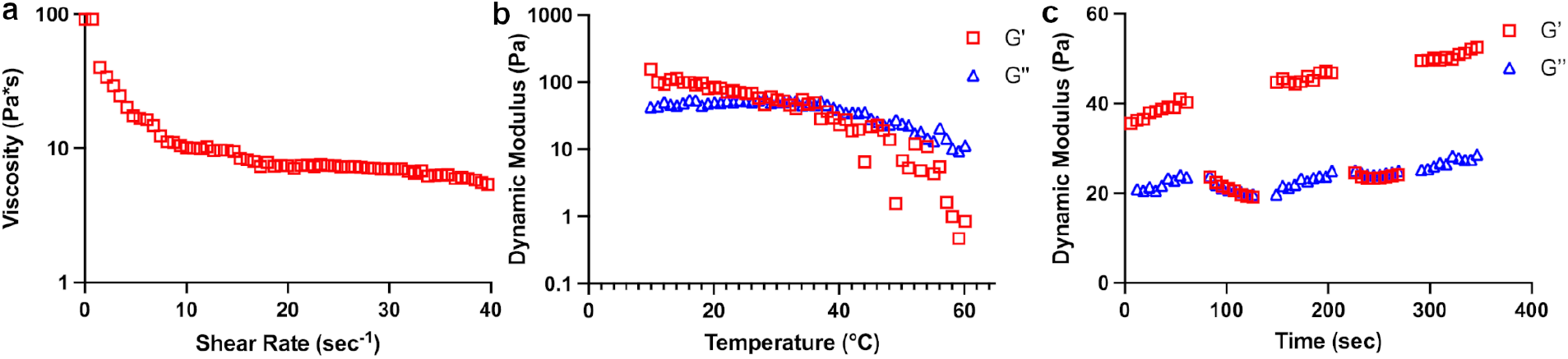
Rheological analysis of the alginate-E3/K3 hydrogels. (a) shear-thinning behavior of the hydrogels with increasing shear rate. (b) Temperature sweep data showing a sharp breakdown of the gels at 45°C, whereafter the structure of the gel was disrupted. (c) The gels show excellent recovery over multiple cycles of low (1%) and high (100%) strain over a period of 350 seconds.

### Rheological Characterization

The steady flow curves of the alginate-E3/K3 gels showed decreasing viscosity with increasing shear rate, indicating shear-thinning behavior (**Figure 4a**). This was further highlighted by fitting a power law model in the viscosity vs shear rate plot.^45^ The gels showed a high flow consistency index of 47.95, demonstrating strong crosslinking and structural integrity and a power law index of -0.6 which confirmed their shear thinning nature.^46^ The gels showed good structural integrity at 25°C as well as at 37°C with a decreasing trend in the storage modulus with increasing temperature. However, at 45°C the storage modulus dropped sharply below the loss modulus indicating loss of structural integrity presumably due to the denaturation of the peptide crosslinks (**Figure 4b**).^47^ Interestingly, the gels maintained their structural integrity and showed significant recovery when cycled through multiple high (100%) and low (1%) strain conditions (**Figure 4c**). This demonstrates that the CC crosslinked gels can be injected under shear without loss of structural integrity, a property useful in many biomedical applications.^48^

### Macro- and Microscale Structure

To further confirm their mechanical suitability for cell encapsulation, we isolated peritoneal MΦs and cultured them on tissue culture plastic (TCP) or encapsulated in alginate crosslinked with Ca^2+^ or CC peptides for 24h. Conventional mechanisms of Ca^2+^ alginate crosslinking were investigated including the use of CaCl_2_ or CaSO_4_ alongside the CC crosslinking system.^11,16,49^ In all encapsulated gels, similar cellular morphology was observed which was distinct from cells cultured on TCP. Notably MΦs on TCP exhibited visible spreading (**Figure S11a**) whereas encapsulated MΦs at the surface and in the bulk of the gel exhibited a rounded morphology (**Figure S11b-g**). We next used scanning electron microscopy (SEM) to characterize microscale pore size of 1% Ca^2+^ (**Figure 5a**) and CC (**Figure 5b**) crosslinked alginate gels. Similar to tradtional Ca2+ crosslinked gels CC crosslinked gels also displayed microporous structure indicative of hydrogelation. The median pore diameter of CC crosslinked gels was 0.596 μm, approximately one fourth that of Ca^2+^ crosslinked gels at 2.42 μm (**Figure 5c**). Pore diameters in Ca^2+^ crosslinked gels exhibited a pronounced positive skew in their distribution (**Figure 5d**) which was largely absent in CC crosslinked gels (**Figure 5e**) which displayed a more tightly controlled distribution of pore diameters. SEM analysis confirmed that, similarly to conventional Ca^2+^ crosslinked alginate gels, CC crosslinked gels form a microporous structure to facilitate cell signalling and nutrient transport.

**Figure 5.**
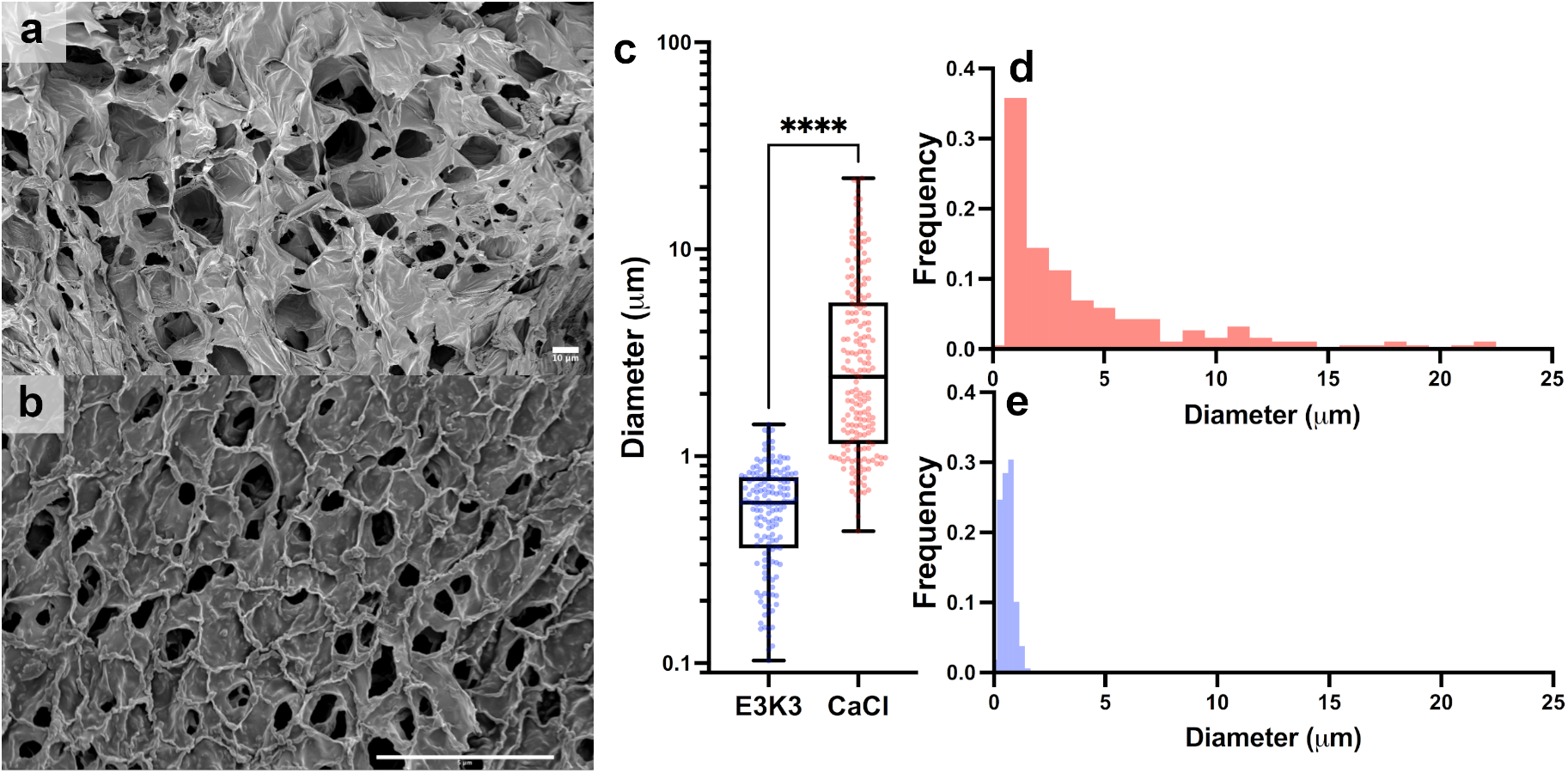
SEM analysis of hydrogel pore size in CC and Ca^2+^ crosslinked alginate hydrogels. (a) Representative SEM micrograph of a CaCl_2_ crosslinked hydrogel. Scale bar 10 μm. (b) Representative SEM micrograph of a CC crosslinked hydrogel. Sclae bar 5μm. (c) Pore diameters of Ca^2+^ and CC crosslinked hydrogels. Statistical comparison performed with student’s t-test. **** P<0.0001. (d) Histogram of pore diameter distribution in Ca^2+^ crosslinked gels. (e) Histogram of pore diameter distribution in CC crosslinked hydrogels.

### Inflammatory properties of Ca^2+^ and CC crosslinked hydrogels

We next confirmed that extracellular Ca^2+^ was the primary driver of IL-1β secretion and that the alginate used was not contaminated with LPS. We measured IL-1β secretion from peritoneal MΦs encapsulated in Ca^2+^ crosslinked gels without exogenous LPS addition in the presence or absence of the LPS-induced-inflammation inhibitor polymyxin-B.^50^ Data showed no significant difference in IL-1β levels in the presence or absence of polymyxin-B, indicating that the alginate used contains negligible LPS (**Figure S12**). We next measured IL-1β release by murine peritoneal MΦs and bone marrow-derived dendritic cells (BMDCs) encapsulated in CC or Ca^2+^ crosslinked hydrogels following LPS stimulation. ELISA data indicated that both MΦs and BMDCs encapsulated in Ca^2+^ crosslinked gels secreted significantly more IL-1β compared to CC crosslinked gels indicating the role of extracellular Ca^2+^ in driving inflammation (**Figure 6a**). This is in agreement with previous studies that demonstrate that extracellualr Ca^2+^ is a secondary messenger in IL-1β secretion in response to LPS insult ^22^. This response was also confirmed in primary human MΦs encapsulated in Ca^2+^ or CC crosslinked alginate gels (**Fig S13**). We further characterizated the phenotype of the encapsulated innate immune cells by degrading the gels with 1mg/mL alginate lyase for 1 hour for flow cytometric analysis. To facilitate this, cells were washed and stained for activation markers CD80, CD40, CD86, MHC-II, and F4/80 (**gating strategy Figure S14**). Data indicated higher expression of CD80 (**Figure 6b**) and CD40 (**Figure 6d**) in Ca^2+^ crosslinked gels compared to CC crosslinked gels. Little to no difference was observed in the expression of MHC-II (**Figure 6c**) across all groups. Interestingly, CD86 expression on MΦs cultured in CC gels was lower than TCP or Ca^2+^ crosslinked gels (**Figure 6e**).

**Figure 6.**
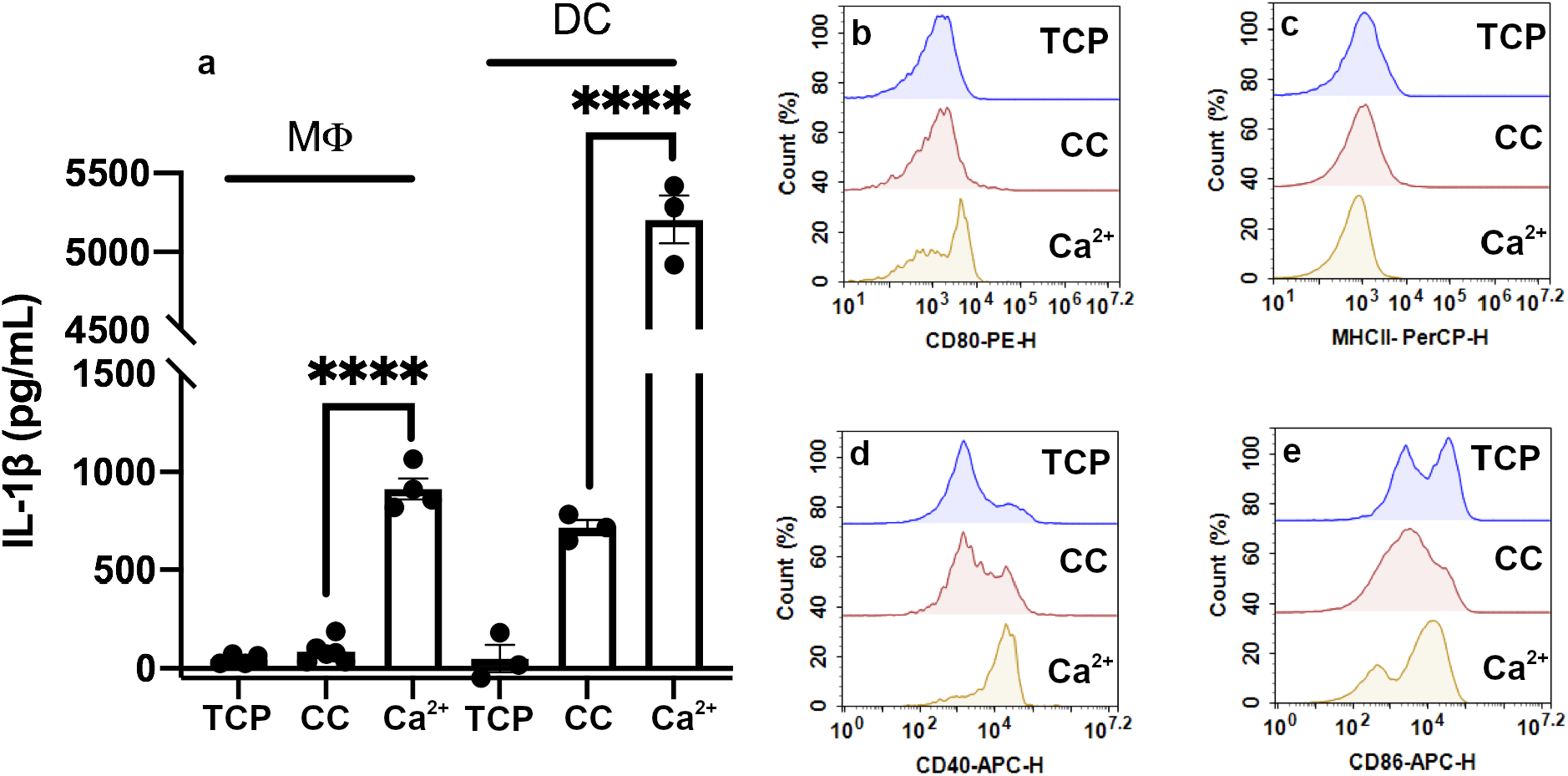
Assessment of the proinflammatory potential of Ca^2+^ and CC alginate gels *in vitro*. (a) Primary MΦs and BMDCs cultured in Ca^2+^ crosslinked alginate gels produce significantly greater titers of IL-1β in response to LPS insult compared to CC crosslinked gels. (b-e) Histograms of MΦ surface expression of CD80 (b), MHC-II (c), CD40 (d) and CD86 (e) under various culture conditions. **** P<0.0001 by student’s t-test

### In Vivo Immunogenicity

To investigate *in vivo* effects of Ca^2+^ or CC crosslinking, we injected mice peritoneally with 150 μL of 0.5% alginate sol or gels crosslinked via Ca^2+^ or CC peptides. Mice were euthanized 24h later and peritoneal lavage was collected and pooled within groups. Cells were washed and stained and analyzed to determine the identity and activation status of infiltrating cells (**gating strategy Figure S15**). The populations of CD11b^+^ (**Figure 7a-d**) and CD11b^-^ (**Figure 7e-h**) cells were found to contain multiple cell types. Within the CD11b^+^ cells, populations of small and large peritoneal MΦs (CD11b^+^ F4/80^hi/int^ Ly6-G^-^),^51^ eosinophil-like cells (CD11b^+^ F4/80^+^ Ly6-G^+^),^52^ and neutrophils (CD11b^+^ F4/80^-^ Ly6-G^+^), were identified (**Figure S16**).^53–56^ Similarly, CD11b^-^ cells were identified as CD11b^-^ neutrophils (CD11b^-^, F4/80^-^, Ly6-G^+^) and heterogenous population of infiltrating cells including DCs (CD11b^-^, F4/80^int/-^, Ly6-G^-^) (**Figure S17**).^55–57^ The heterogenous triple negative population (highlighted in red: Figure 7e-h) was used to determine an appropriate threshold for MHC-II activation, as it likely represented both MHC-II expressing DCs and other MHC-II non-expressing leukocytes. This population displayed a characteristically bimodal MHC-II response (**Figure 7m-p**). The point of demarcation was used to differentiate between MHC-II^+^ and MHC-II^-^ expression. MHC II expression was used to determine the degree of activation of peritoneal MΦs (**Figure 7i-l**) and was found to be lower in mice treated with CC crosslinked alginate (31.8%) compared to Ca^2+^ crosslinked alginate (78.2%). Interestingly, activation of MΦs was also observed in mice treated with alginate sol. Notably, these findings are in agreement with our *in vitro* observations that CC crosslinked gels are less inflammatory compared ton Ca^2+^ crosslinked gels (**Figure 6e**). Beyond the activation state of MΦs, the identity of cells recovered from the peritoneum was substantially different in the CC crosslinked condition, with a major increase in the proportion of granulocytes (**Figure 7d**). In particular, eosinophils (6-fold) and CD11b^+^ neutrophils (3-fold) were elevated compared alginate sol or Ca^2+^ crosslinked gels. This may indicate that the immune response to the CC crosslinked gels *in vivo* is much more complex than MΦ activation state and that the recruitment of granulocytes may be a valuable avenue for future study of CC crosslinked hydrogels. The major increase in eosinophils in particular may be relevant to tissue engineering due to their adjunct role in promoting wound healing.^58,59^

**Figure 7.**
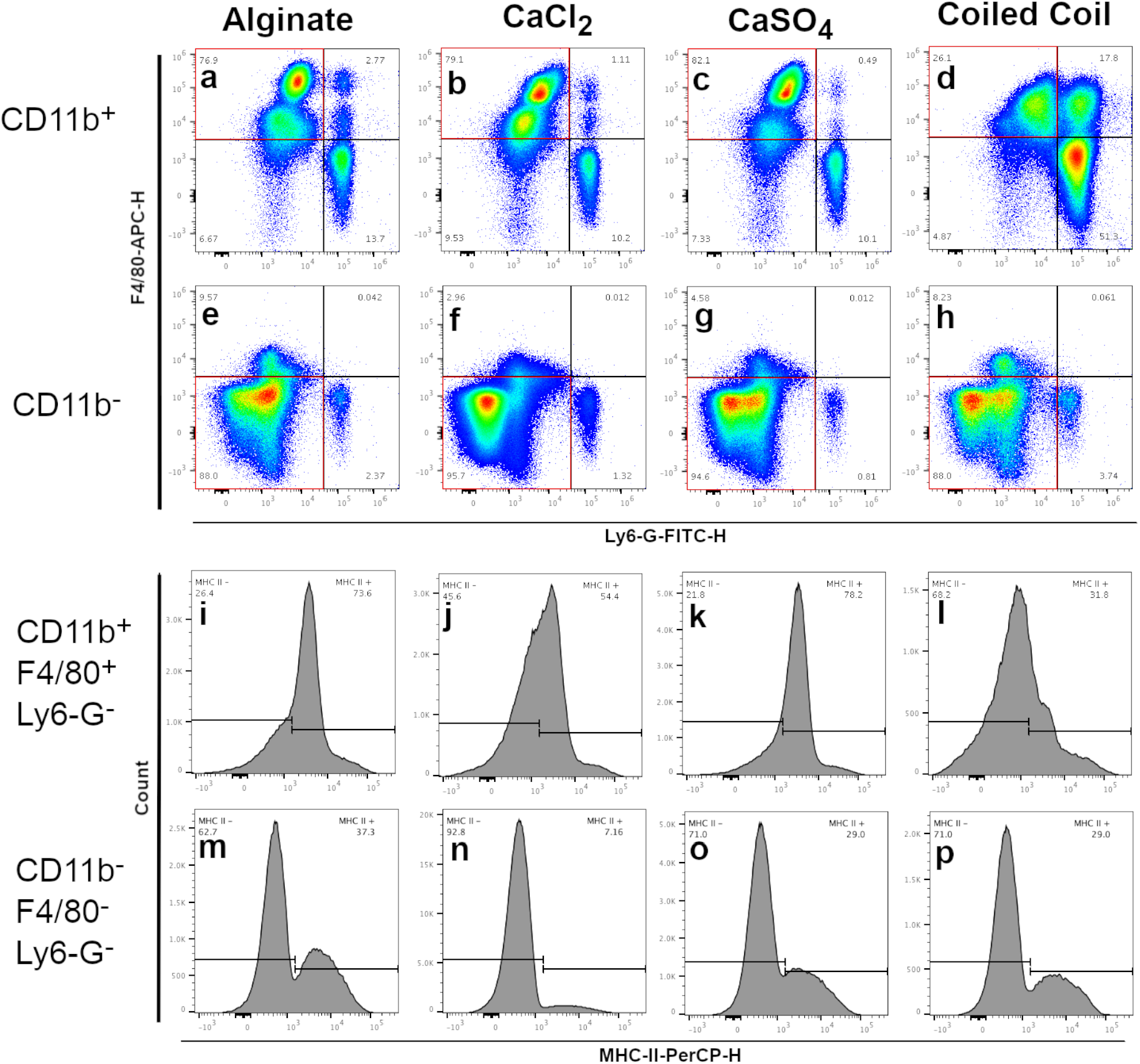
Flow cytometric analysis of peritoneal lavage from mice injected with CC or Ca^2+^ crosslinked alginate. (a-d) CD11b^+^ cells from all groups included a substantial F4/80^+^ Ly6-G^-^ population corresponding to macrophages, as well as well as a F4/80^-^ Ly6-G^+^ population corresponding to neutrophils. Notably the CC group also exhibited a substantial CD11b^+^, F4/80^+^, Ly6-G^+^ triple positive population corresponding to eosinophils (d). (e-h) CD11b^-^ cells in all groups primarily exhibited a CD11b^-^, F4/80^-^, Ly6-G^-^ population which likely includes dendritic cells among other leukocytes as well as a CD11b^-^, F4/80^-^, Ly6-G^+^ population corresponding to CD11b^-^ neutrophils. (i-l) Among macrophage-like cells the CC group (l) exhibited the least activated phenotype of MHC-II expression. (m-p) Among CD11b^-^, F4/80^-^, Ly6-G^-^ triple negative cells all groups exhibited a bimodal MHC-II profile, useful for determining an MHC-II gate.

Despite the many advantages alginate hydrogels offer as cell scaffolds, one notable drawback is the inflammatory nature of extracellular Ca^2+^ which can limit the use of conventionally crosslinked alginate gels in the presence of immune cells.^21^ Grafting heterodimeric coiled coil-forming motifs onto alginate offers a mechanism for Ca^2+^-free mechanically reversible crosslinking in alginate through supramolecular self-assembly which can greatly reduce MΦ-driven inflammation compared to conventional Ca^2+^ ion crosslinking methods. Coiled coils are known to display a tunable sequence-structure-function relationship making them a versatile motif for polymer crosslinking.^60^ Alterations to the heptad structure of designed coiled coil forming peptides can be used to tune the melting point and pH sensitivity of the supramolecular assembly which may find use in the design of stimulus responsive cell scaffolds.^31,61^ Notably, the use of coiled-coil forming peptides to crosslink hydrogels is not unique to alginate, but may find use crosslinking a wide range of natural polymer cell scaffolds.

## Conclusion

Here we describe synthesis of the graft copolymers: alginate-E3 and alginate-K3, the first to conjugate CC peptides to alginate for study at the macroscale for cell encapsulation. We demonstrate that alginate-E3/K3 can be crosslinked in the absence of Ca^2+^ and that they significantly reduce IL-1β production from encapsulated primary MΦs in response to an LPS insult *in vitro* compared to Ca^2+^ crosslinked gels making them more optimal for tissue engineering applications to avoid potential development of non-healing wounds. In vivo, the CC crosslinked gels were also found to be minimally inflammatory. We also observed a marked increase in granulocytes, particularly eosinophils, in the peritoneal space of mice injected with the CC crosslinked gels. Further crosslinking via supramolecular CCs can be extended to other polymers for fabricating functional hydrogels.

## SUPPORTING INFORMATION

Supporting Information includes: Expanded detail on methods used, TableS1: sequence charge and mass of E3 and K3 peptides. Scheme S1: Solid phase peptide synthesis procedure. Figures S1-S3: HPLC traces of purification and background signal. Figures S4, S5 MALDI mass spectra of E3 and K3. Figures S6-S8 ^1^H NMR spectra for maleimidoethylamine, alginate and BMPH-alginate. Figure S9: CC crosslinked alginate gels are self supporting when inverted. Figure S10: Circular dichroism spectroscopy background trace for both water and alginate. Figure S11: bright field microscopy of primary MΦ encapsulated in CC and Ca^2+^ crosslinked gels at the well surface and in the bulk of the gel. Figure S12: IL-1β secretion of primary MΦ cultured in alginate sol the presence or absence of polymyxin B. Figure S13: IL-1β secretion of human PBMC derived MΦ cultured on TCP or in CC or Ca^2+^ crosslinked alginate. Figure S14, S15: Gating strategies for flow cytometric analysis of cells exposed to CC or Ca^2+^ crosslinked gels *in vitro* and *in vivo*.

## ACKNOWLEDGEMENTS

The authors would like to thank the High-resolution NMR Facility and Deaprtment of Chemistry at Washington University in St. Louis for their assistance with NMR spectroscopy. The authors would like to thank the Washington University Center for Cellular Imaging and the Medical School at Washington University for assistance with scanning electron microscopy.

## Author Contributions

Z.C. and J.S.R designed and executed experiments, interpreted data and prepared the manuscript. C.O. performed CD spectroscopy. G.L and A.G performed rheology. P.S and M.F. assisted with flow cytometry. D.D.K. purified peptides with HPLC. All authors have given approval to the final version of the manuscript.

## Notes

The authors declare no competing financial interests.

## Table of Contents Graphic

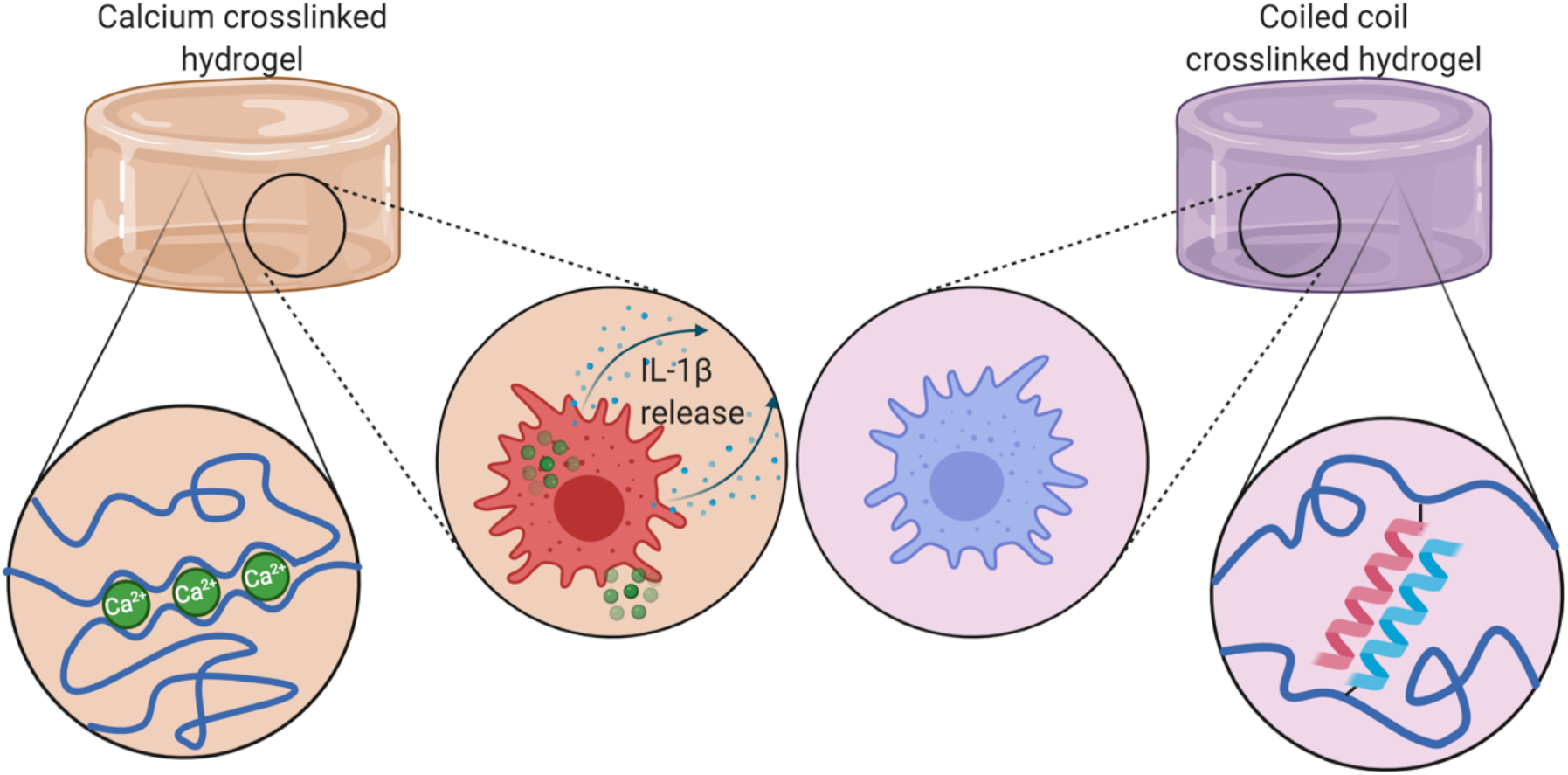

