## Supplemental Methods and Figures for "Coiled Coil Crosslinked Alginate Hydrogels Dampen Macrophage-Driven Inflammation"

**Additional Experimental Details**:

*Solid Phase Peptide Synthesis*:

Peptides were synthesized using a LibertyBlue automated peptide synthesizer (CEM, Matthews NC) by solid phase peptide synthesis following manufacturer's instructions. Reaction Scheme: (**Figure S1**). Peptides were synthesized using preloaded Wang resin (ThermoFisher, Waltham, MA(ThermoFisher)) and were not N-terminally acetylated. Fmoc-amino acids were purchased from Chem-Impex (Wood Dale, IL). First, resin was swelled in N,N-Dimethylformamide (DMF) (Arcos Organics) for 2 minutes then each amino acid was added sequentially. Before each amino acid was added, the FMOC protecting group was cleaved with 20% piperidine in DMF for 2 minutes with microwave heating to 90 °C and bubbling with $N_2$ then drained. The resin was then washed 3 times with 5 mL of DMF to remove residual piperidine. Amino acids were added by suspending the resin in a cocktail composed of 3 mL of 0.2M FMOC-protected amino acid in DMF, 1mL of 1 M Oxyma-pure in DMF and 1 mL of 1 M diisopropylacrbodiimide in DMF. The coupling cocktail was heated to 90 °C and bubbled with $N_2$ for 2 minutes before draining. This process was repeated for each amino acid synthesizing the peptide from the C to N terminus. Once the desired sequence was completed, the resin was cleaved one more time as described to remove the N-terminal FMOC group and washed with several volumes of dichloromethane. The resin was dried with $N_2$ then under vacuum for 30 minutes. At this point the peptide conjugated resin was either immediately cleaved or stored at -20 °C for future cleavage. Peptides were cleaved from resin using a cleavage cocktail composed of a 92.5 : 2.5 : 2.5 : 2.5 volumetric ratio trifluoroacetic acid : deionized water : triisopropylsilane : 1,2-ethanedithiol (EDT). Resin was mixed on an orbital shaker with least 10 mL of cleavage cocktail per gram of resin for 2 hours. After cleavage, the cocktail was removed using rotary evaporation and the isolated peptide was resuspended in a 3 : 1 volumetric mixture of deionized water : acetonitrile. The suspension was frozen at -80 °C overnight and then lyophilized completely using a Labconco FreeZone 4.5 L - 105C PTFE-coated freeze dryer.

*High Performance Liquid Chromatography (HPLC)*:

Peptide purification by reverse-phase HPLC was conducted using an UltiMate 3000 UHPLC system (ThermoFisher) Using a Zorbax reverse phase PrepHT C18 column (Agilent Technologies, Santa Clara, CA) using a constant speed ramp procedure. Ultrapure water containing 0.5% v/v trifluoroacetic acid (TFA) (Sigma-Aldrich, St. Louis, MO (Sigma)) was used as the aqueous phase and acetonitrile (Sigma) containing 0.5% v/v TFA was used as the organic phase.

*Alginate-Peptide Conjugation*:

Alginate-peptide conjugates were formed using a 2 step process. first N-β-maleimidopropionic acid hydrazide (BMPH) (Thermo) was conjugated to alginate by carbodiimide reaction. To do this, first an MES buffer was created by dissolving 2-(N-morpholino)ethanesulfonic acid (MES) (Sigma) in Deionized water to a final concentration of 0.5M and pH adjusted to 5.0. Sodium alginate (Thermo) was then dissolved completely in MES buffer to a concentration of 1% w/v under stirring. A twofold molar excess to the desired final degree of substitution of BMPH was added to the solution (e.g. for a theoretical 2% final substitution of peptide to alginate BMPH was added to a molar ratio of 0.04 mol[BMPH]/mol[alginate monomer] ). At the same time 1-ethyl-3-(3-dimethylaminopropyl) carbodiimide (EDC) (G-Biosciences, Overland, MO) was added in eightfold molar excess to the final degree of substitution (e.g. for a theoretical 2% final substitution of peptide to alginate EDC was added to a molar ratio of 0.16 mol[EDC]/mol[alginate monomer]). This solution was allowed to react overnight stirring at room temperature. The Alginate-BMPH was recovered by precipitation with a 4 fold volume excess of ethanol added to the stirring solution. The obtained precipitate was collected and dried by vacuum filtration on a piece of No. 1 Whatman paper in a Buchner funnel. To ensure complete removal of excess MES from the obtained alginate, the product was again dissolved in deionized water and precipitated with ethanol and dried with the same method as previous. Once largely dried by vacuum filtration the alginate-BMPH was completely dried under vacuum overnight. At this point the Alginate-BMPH could be safely stored and analyzed, for example, by NMR spectroscopy. The second step in creating the alginate-peptide conjugate molecule was addition of peptide to alginate-BMPH through maleimide thiol crosslinking with cysteine residues on E3 and K3. First PBS was pH adjusted to 7.2. Alginate was dissolved in the PBS to a concentration of 0.5% w/v under stirring. The peptide of interest was the dissolved in the solution at the desired level of substitution (e.g. for a theoretical 2% final substitution of peptide to alginate the peptide was added to a molar ratio of 0.02 mol[peptide]/mol[alginate monomer] ) and the reaction was let stir at room temperature overnight. Once the reaction was completed any free maleimide moieties were quenched with an excess of free cysteine for 30 minutes. Peptide conjugated alginate was collected through vacuum filtration and precipitation through the same method as previously described then redissolved in deionized water at 0.5% w/v and dialyzed using 3500 Da float-a-lyzer cassettes following manufacturer's instructions (Spectrum Chemical, Gardena, CA) against deionized water for 2 days, changing the dialysis bath twice daily. Once dialysis was complete the dialyzed solutions were frozen at -80 C and lyophilized.

*NMR Spectroscopy*:

NMR samples were prepared by dissolving the compound of interest in $D_2O$ spiked with TMS at 1 mg/mL. The samples prepared were alginate, 2% substitution BMPH-alginate, and maleimidoethylamine, in order to distinguish the characteristic peak of a maleimide group. $^1$H

NMR spectra were obtained at 600 MHz using an Agilent DD2 600 MHz NMR spectrometer. NMR spectra were analyzed using MestreNova software.

*FAM-Thiol Fluorometric Conjugation Quantification*:

To quantify the final degree of substitution a reaction was performed using the same methods as described in *alginate-peptide conjugation* for binding peptide to alginate-BMPH, replacing the peptide with FAM-thiol. The reaction was otherwise conducted identically to the one described previously. The conjugation was assessed by fluorescence assay. The alginate-FAM conjugate was dissolved in deionized water at concentrations corresponding to 289.2, 144.6 and 72.3 μM of alginate monomer, the fluorescence of the samples was measured using a Synergy H1 microplate reader (BioTek, Winooski, VT) and compared to a standard curve with excitation at 493 nm and emission measured at 517 nm to obtain the resulting molar concentration of FAM. From there the molar ratio of FAM to alginate monomer was computed and for each concentration of alginate-FAM and compared to ensure consistency.

*Circular Dichroism Spectroscopy*:

CD spectra were recorded on a Jasco J-810 circular dichroism spectrometer. Solutions were prepared in PBS at either 0.27 mg/mL (E3, K3, and E3+K3) or 1.0 mg/mL (Alg-E3, Alg-K3, Alg-E3+Alg-K3). The spectra consist of 3 accumulations collected at 20°C in a 1 mm quartz cuvette between 300 and 190 nm at 50 nm/min with 1 nm bandwidth and 0.1 nm data pitch. The background was subtracted and the raw values were converted to molar ellipticity manually.

*Hydrogel Formation*:

CC crosslinked hydrogels were formed by dissolving alginate-E3 and alginate-K3 at the desired final concentration (i.e. for a 1% final concentration both a 1% solution of alginate-E3 and alginate-K3 were prepared) in 1x PBS by heating to 45 ˚C and intermittently vortexing. Once both polymers were completely dissolved equal volumes of alginate-E3 and alginate-K3 were mixed by pipetting up and down and allowed to set at room temperature for 15 minutes. If cells were to be encapsulated in the gels, then cells were pelleted and resuspended in the alginate-E3 solution to achieve a final cell concentration of $10^6$ cells/mL (i.e. at a concentration of $2\times10^6$ cells/mL in the alginate-E3 solution). $CaSO_4$ crosslinked gels were formed using similar methods to those previously described [22]. In short, 375 µL of 1.3% alginate solution in which cells were resuspended to a concentration of $1.3\times10^6$ cells/mL was drawn into a 1 mL syringe and 125 µL of a 183 mM slurry of $CaSO_4$ was drawn into the tip of a 1000 µL pipette. The pipette tip was then inserted into the tip of the 1 mL syringe and the two solutions were quickly mixed back and forth approximately 5 times and then extruded into 96-well plate at a volume of approximately 100 µL per well. To form $CaCl_2$ crosslinked gels, cells were suspended at a concentration of $1.3\times 10^6$ cells/mL in a 1.3% solution of alginate. 75 µl of the solution was pipetted into the wells of a 96-well plate. Then to gel, 25 µL of a 183 mM $CaCl_2$ solution was added by pipetting directly into the solution. For culture, the gels were covered in 100 µL of RPMI-1640 media supplemented with 10% fetal bovine serum and 1% penicillin/streptomycin.

*Rheology*:

Shear thinning properties of the alginate gels crosslinked by E3/K3 peptides was studied using Discovery HR-2 rheometer. Sample solutions were loaded between a parallel plate geometry of 8mm diameter. A shear rate sweep of 0-40 $s^{-1}$ at 25°C was applied. Temperature sweeps were performed from 10-60 °C at 1% strain and an angular frequency of 10 rad/s. Recovery of alginate gels was studied by monitoring storage modulus changes at low (1%) and high (100%) strain values. All analysis was performed using TRIOS software and viscosity values with respect to shear rate were fit in a power law model.

*Primary MΦ Isolation*:

Primary MΦ were isolated using a similar protocol as described previously [35]. Briefly, female C57BL6/J mice were euthanized by isoflurane inhalation followed by rapid cervical dislocation. Immediately after euthanasia the peritoneal space was lavaged with 10 mL of sterile PBS. After PBS was injected the mice were agitated for 30 seconds to dislodge peritoneal cells. After agitation, 8 mL of peritoneal fluid was extracted, taking special care to avoid organ or blood vessel puncture. The extracted fluid was centrifuged at 300 xg for 5 minutes at 4 °C to pellet cells. Cells were then resuspended in 5 mL of RPMI-1640 media supplemented with 10% fetal bovine serum and 1% penicillin/streptomycin and counted. Peritoneal MΦ were either stained for flow cytometric analysis or used for study immediately after isolation.

*DC Isolation and Expansion*:

Under sterile conditions, murine bone marrow was isolated from femurs and tibias of female C57B6 mice. The gathered bone marrow was treated with RBC lysis buffer for two minutes and washed with saline buffer twice. Washed monocytes were resuspended in complete growth media and were cultured for six days supplemented with GM-CSF and IL-4 to promoted differentiation into dendritic cells. The growth media were replenished on day 3 and day 5, and on day 6 the cells were collected and purified using EasySep Mouse $CD11c^+$ Selection Kit following the manufacturer's instructions (Stemcell Technologies, Vancouver, BC, Canada).

*Determination of Cell Response to CC and $Ca^{2+}$ Encapsulation In Vitro*:

Cells were isolated as described above and encapsulated in hydrogels as described previously. After encapsulation cells were allowed to rest for 15 minutes. After that time period cells were stimulated with lipopolysaccharide to a final concentration of 100 ng/well (500 ng/mL). 24 hours after LPS stimulation supernatant was collected for later analysis with ELISA. For analysis of cells, media was aspirated and replaced with a solution of 1 mg/mL alginate lyase (Sigma). Gels were then incubated with alginate lyase at 37 °C for 1 hour, collected, washed and stained for flow cytometric analysis.

*Light Microscopy*:

Cells were imaged in gels using light microscopy with a Lionheart automated microscope (BioTek, Winooski, VT) using a 20x magnification objective.

*Scanning Electron Microscopy (SEM)*:

Samples were prepared by first forming gels at 1% w/v crosslinked either with $CaCl_2$ of CCs then flash freezing them by submersion in excess liquid nitrogen. The gels were immediately lyophilized to remove all water. Samples were sputter coated with gold using a Leica EM ACE600 Sputter Coater. Samples were imaged using a Thermo Fisher Helios 5 UX Focused Ion Beam Scanning Electron Microscope. Each type of gel was imaged in duplicate, using two separately prepared samples.

*Image Analysis*:

Images were analyzed manually using FIJI image processing software. For measurement of pore diameter the approximate longest and shortest axis were measured and the diameter was determined to be the average of those measurements.

*Enzyme Linked Immunosorbent Assay (ELISA)*:

Supernatants collected for ELISA analysis were analyzed using an invitrogen mouse IL-1β uncoated ELISA kit (Thermo) following manufacturer's instructions. Briefly, wells of a high binding clear 96-well plate were coated for overnight with the capture antibody prepared as described in the kit. Wells were washed 3x by 1x PBS-t (0.1% Tween-20 in PBS) and dried completely. Then the wells were blocked for 1 hour with a solution of bovine serum albumin prepared as directed by the kit and the wells were again washed 3x with PBS-t. Then either standard solutions or supernatant diluted 10x was added to the wells for 2 hours. The wells were again aspirated and washed 3x with PBS-t. The secondary antibody was prepared as per the manufacturer's instructions and added to wells for 1 hour. The wells were again aspirated and washed 3x with PBS-t. A solution of streptavidin-horseradish peroxidase was prepared per the manufacturer's instructions and added to the wells for 30 minutes. The wells were then aspirated and washed 5x with PBS-t. A solution of 0.4 mg/mL 3,3',5,5' Tetramethyl benzidine in water was added to the wells and allowed to react for 15 minutes at which point the reaction was terminated by addition of an equal volume of 1M phosphoric acid. The absorbance of the wells was measured at 450 nm using a BioTek Synergy H1 microplate reader.

*Flow Cytometry*:

For analysis of *in vitro* cell response cells were isolated from gels by first degrading gels with 1 mg/mL alginate lyase for 1 hour at 37 °C, then washing and staining the cells as previously described [36]. After cells were treated, nonspecific binding sites were blocked with mouse Fc-block solution (BD Biosciences, Franklin Lakes, NJ, USA) in FACS buffer (10% FBS in 1x PBS) on ice. Cells were washed with FACS buffer and divided into two groups. One group was stained with anti-F4/80-APC-eFluor780 (eBioscience), anti-CD80-PE (eBioscience), anti-MHC-II-PerCP (eBioscience) and anti-CD40-APC (eBioscience) while another group was stained with anti-CD86-APC (eBioscience). Both groups were allowed to stain protected from light for 30 minutes. Fluorescence was measured with a Acea Novocyte 3000 flow cytometer (Acea Biosciences, San Diego, CA, USA), with compensation performed using Invitrogen Ultracomp ebeads (Thermo). Flow cytometry data was analyzed on F4/80$^+$ cells gated based on forward and side scatter to exclude debris & doublets further to check the degree of MHC-II, CD40, CD80 and CD86 expression. (**Figure S14**).

For analysis of cells recovered from animals from *in vivo* experiments cells were first obtained through the method outlined previously in *Primary MΦ Isolation*. Once obtained, cells were stained with DAPI (Thermo). Once stained with DAPI, cells were fixed using 4% Paraformaldehyde. After blocking with mouse Fc-Block solution cells were stained with anti-CD11b-PE (eBioscience), anti-F4/80-APC (BioLegend), anti-Ly6-G-AlexaFluor488 (eBioscience) and anti-MHC-II-PerCP (eBioscience) on ice and allowed to stain for 30 minutes protected from light. Fluorescence was measured using the same technique as for cells cultured *in vitro*. Flow Cytometry data was gated on forward and side scatter to exclude debris and doublets, then secondarily gated on DAPI negative cells. These cells were then separated into CD11b positive and negative groups which were each subdivided into 4 groups based on Ly6-G and F4/80 expression. Finally, the level of MHC-II expression in $CD11b^+$, $F4/80^+$, $Ly6\text{-}G^-$ and $CD11b^-$, $F4/80^-$, $Ly6\text{-}G^-$ populations was quantified (**Figure S15**). Flow cytometry data was analyzed using FlowJo software.

*Animal Study*:

Female C57BL/6J mice at 6 weeks of age were obtained from Jackson Laboratory (Bar Harbor, ME, USA). The animals were provided food and water ad libitum and housed in a temperature-controlled environment with 12 hour light and dark cycles. All animal work was approved by the Institutional Animal Care and Use Committee at Washington Univeristy in St. Louis. Mice were separated into 4 groups corresponding to 4 treatments: uncrosslinked alginate, $CaCl_2$ crosslinked alginate, $CaSO_4$ crosslinked alginate and CC crosslinked alginate. Mice were injected intraperitoneally with 150 μL of the appropriate material at 1% w/v and 24 hours later euthanized by isoflurane inhalation followed by cervical dislocation. Immediately after euthanasia, peritoneal cells were collected using the same procedure as described for primary MΦ isolation and stained for flow cytometry as previously described. All efforts were made to minimize animal suffering.

**Supplemental Tables, Figures & Schemes**:

Table S1. Sequences, net charges and masses of E3 and K3 peptides

| Name | Sequence | Net Charge | Mass (Da) |
|---|---|---|---|
| E3 | CGEIAALEKEIAALEKEIAALEKYG | -3 | 2663.05 |
| K3 | CGKIAALKEKIAALKEKIAALKEYG | +3 | 2660.23 |

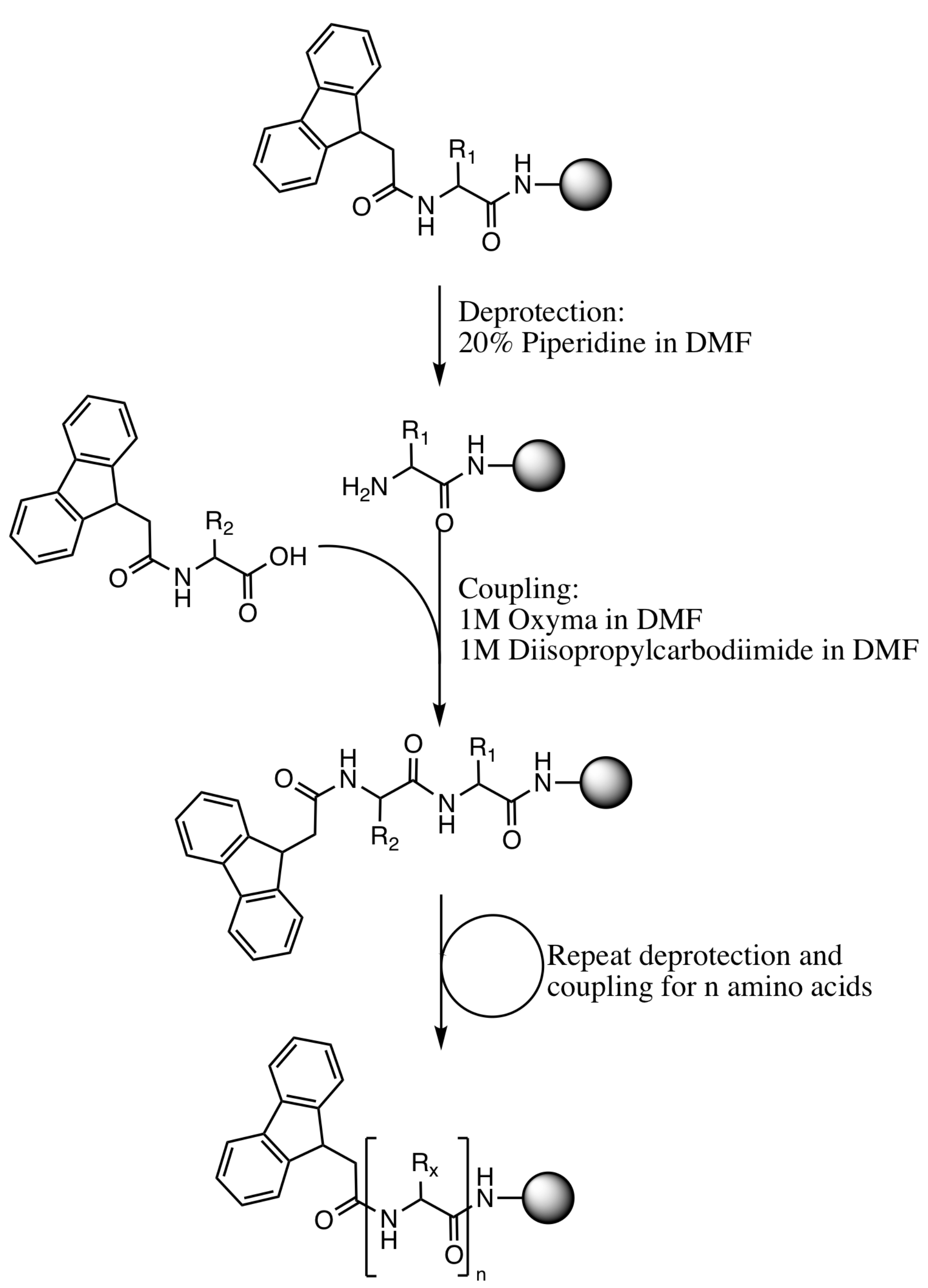


Scheme S1. Schematic of the reaction used for Solid phase peptide synthesis

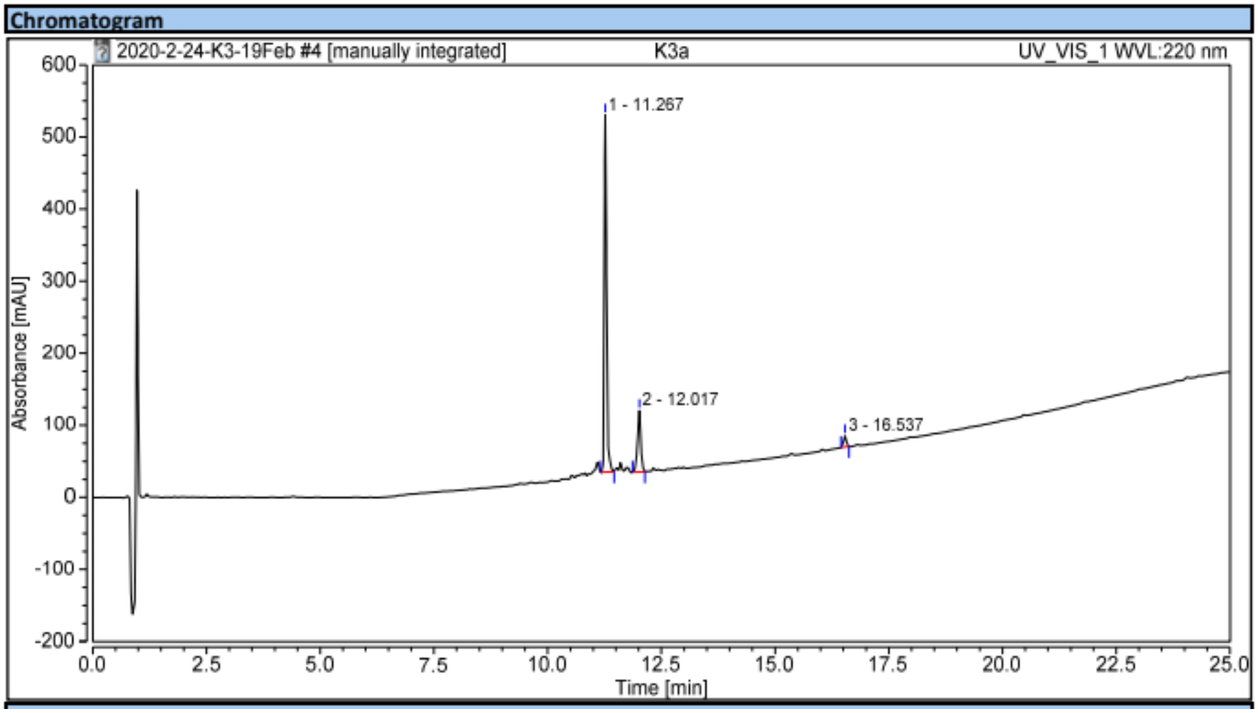


Figure S1. HPLC trace of K3

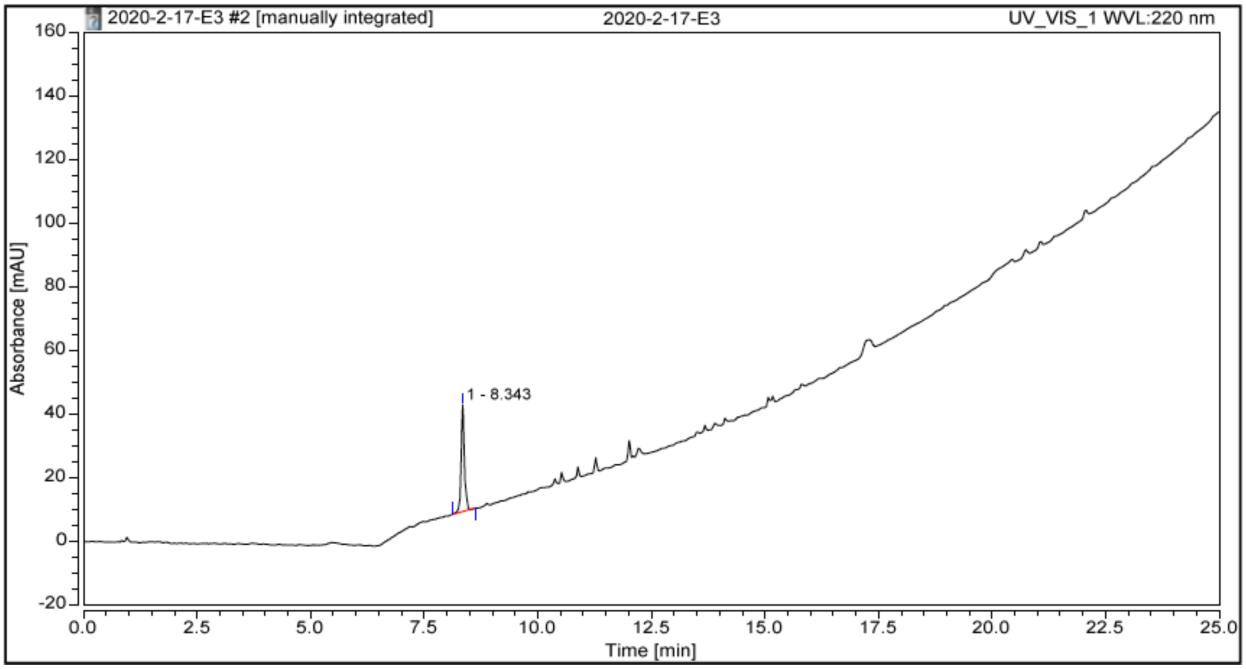


Figure S2. HPLC trace of E3

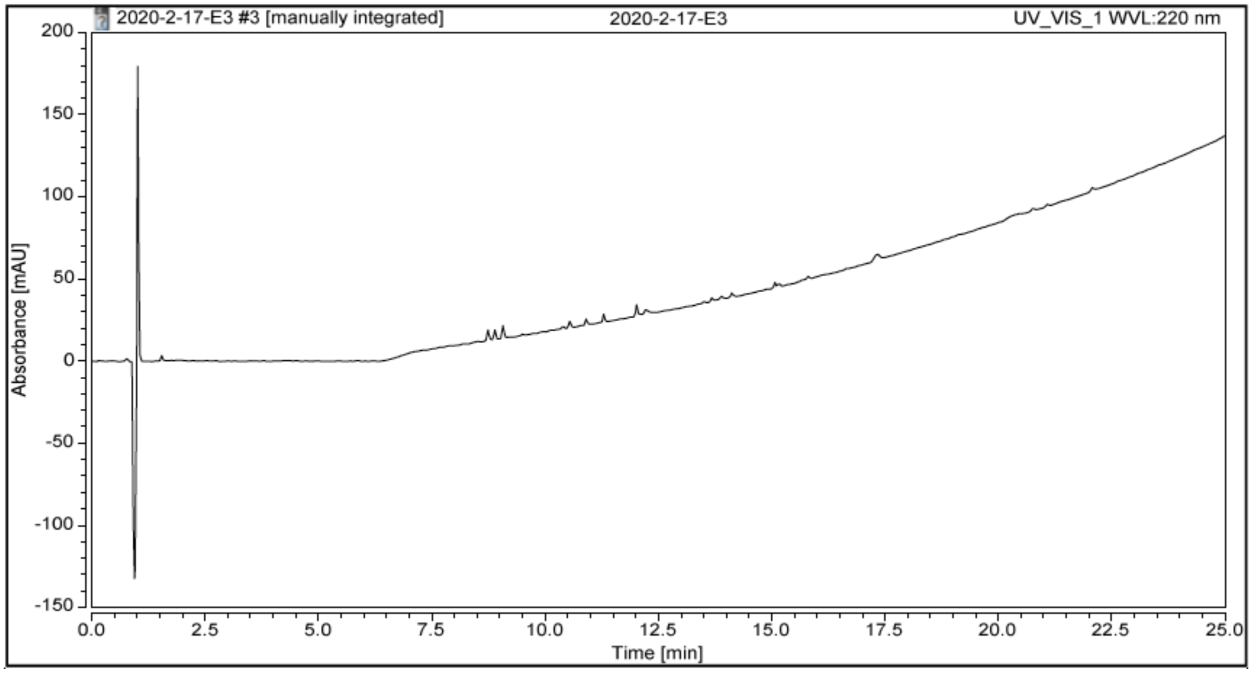


Figure S3. Background HPLC trace

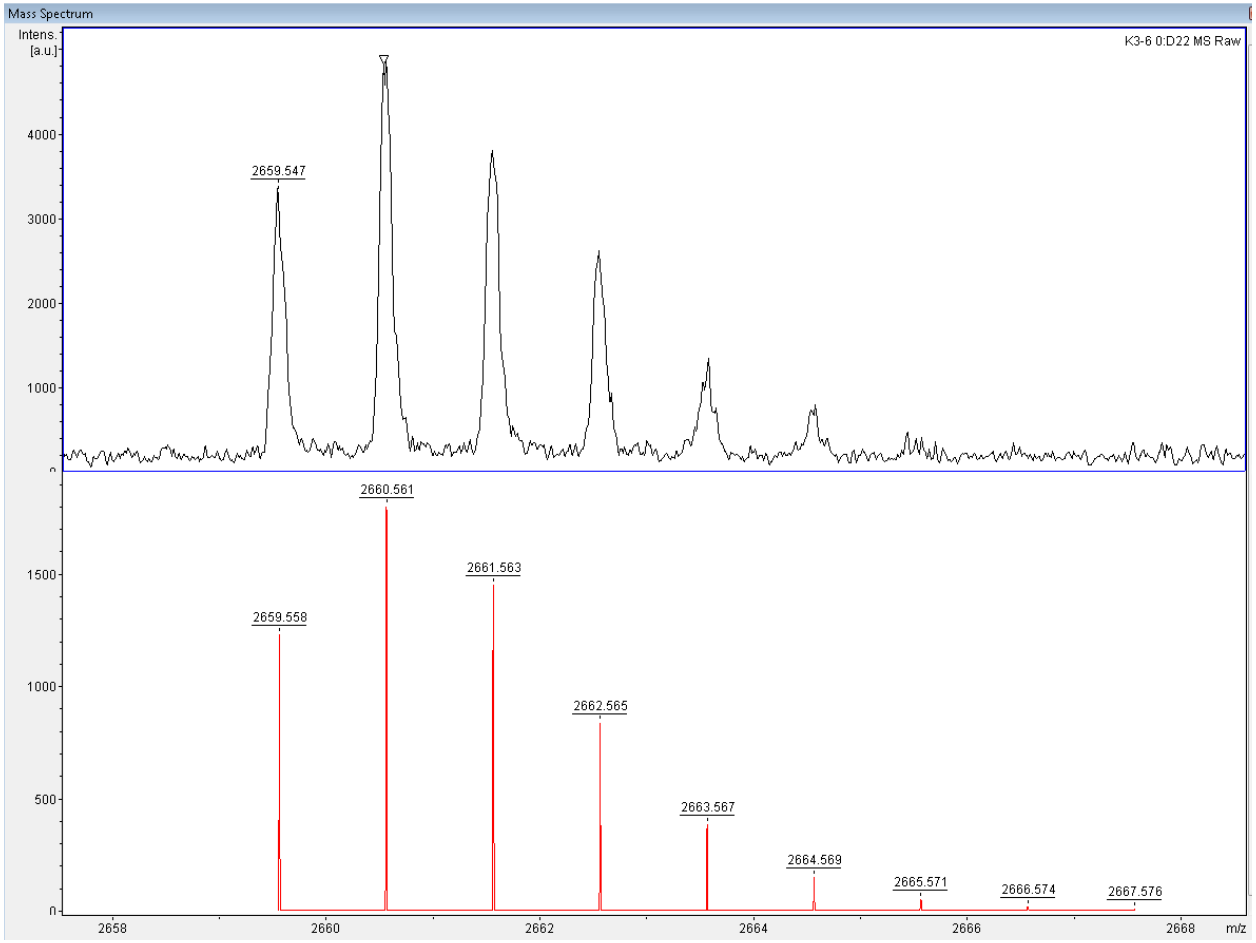

**Figure S4** MALDI mass spectroscopy trace of isolated peptide K3 (measured top, simulated bottom)

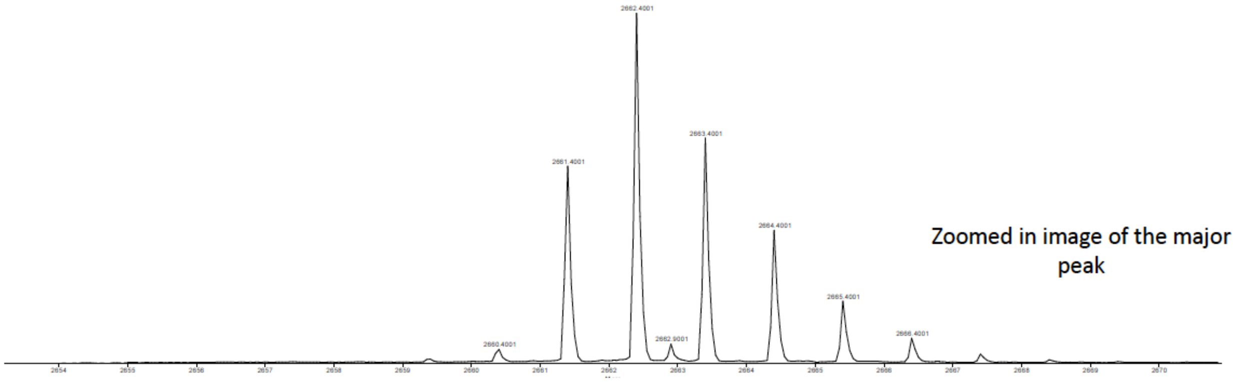


**Figure S5** MALDI mass spectrum of isolated E3 peptide

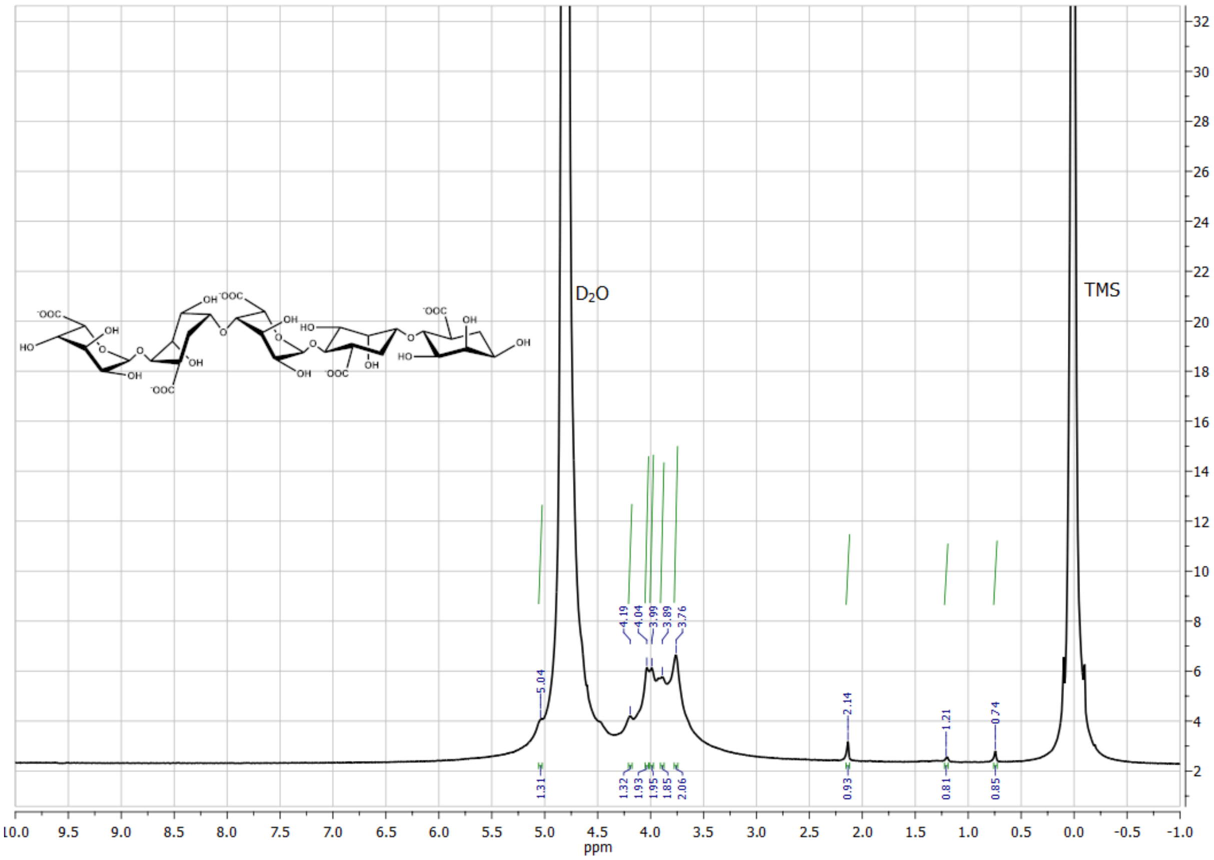


**Figure S6**: $^{1}H$ NMR spectrum of alginate

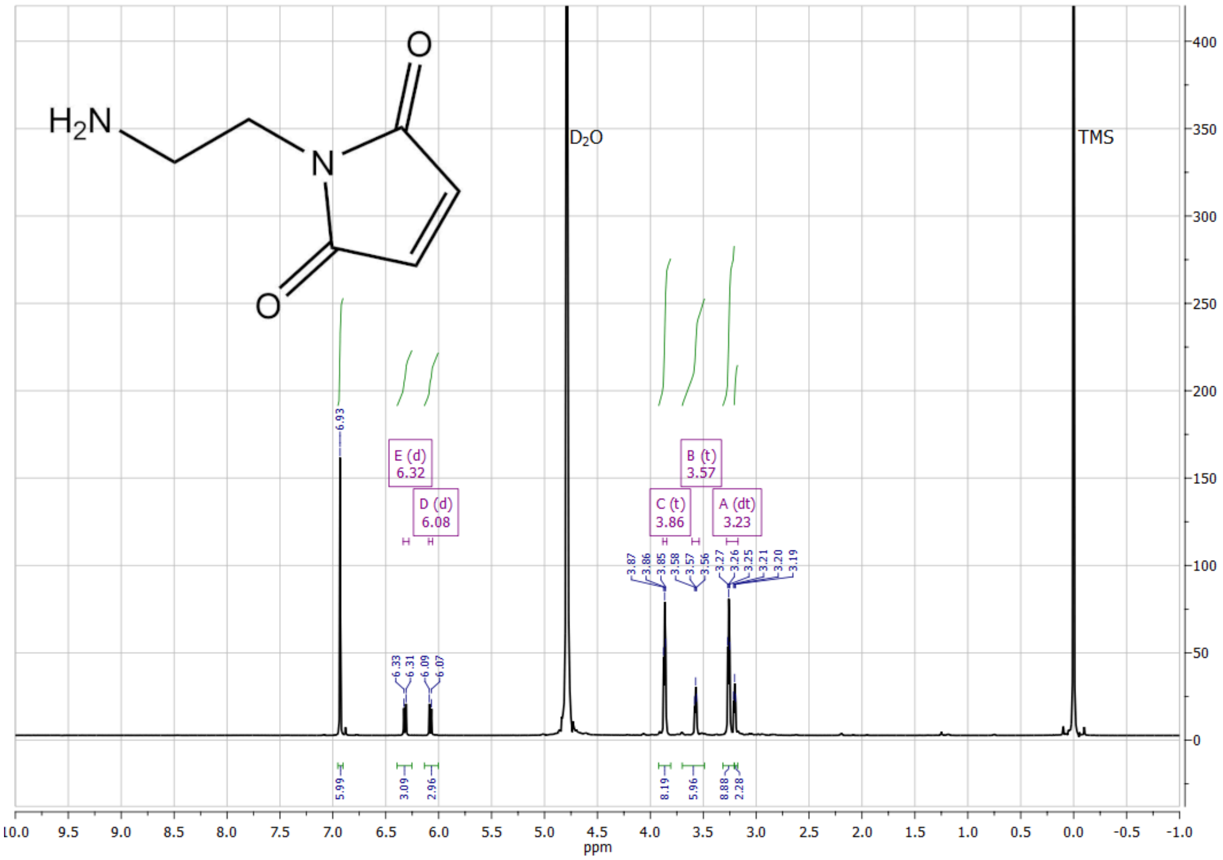


**Figure S7**: [1]H NMR spectrum of maleimideoethylamine

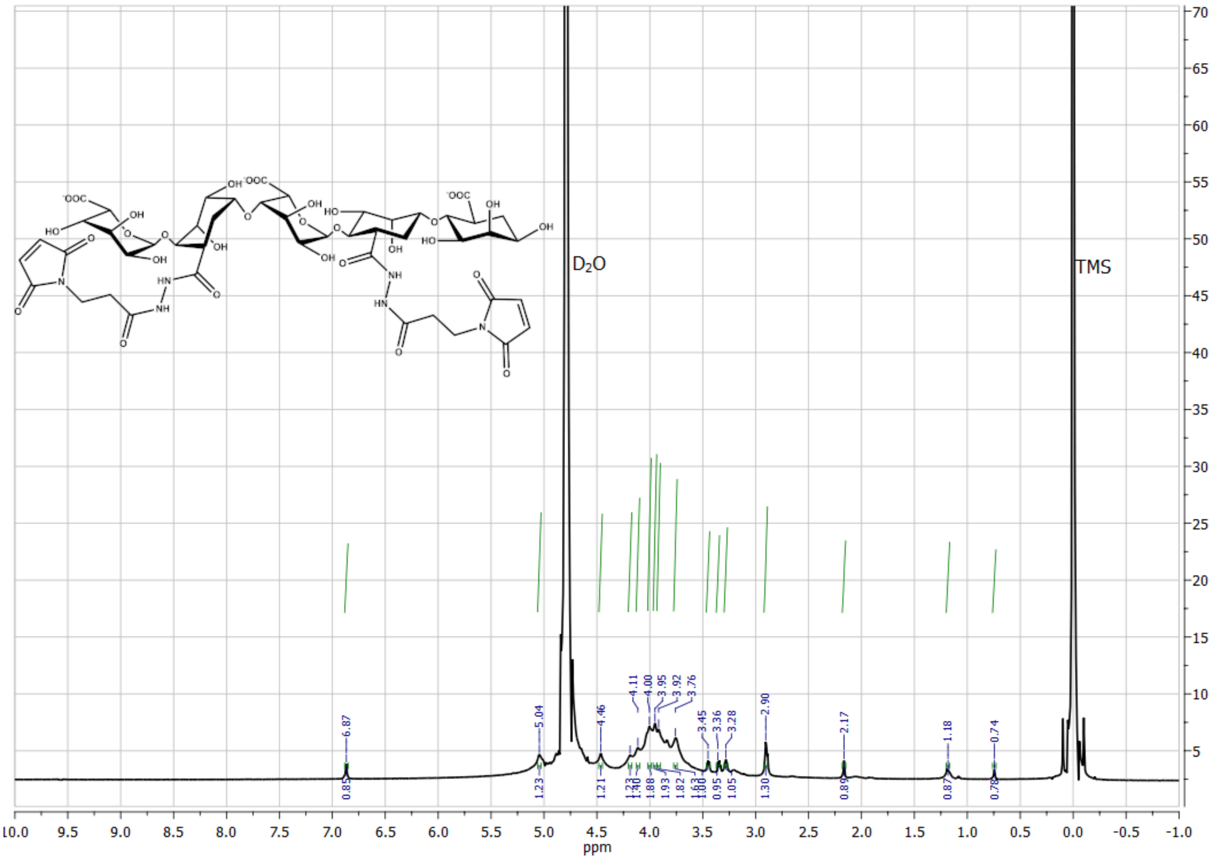


**Figure S8**: $^1$H NMR Spectrum of BMPH-conjugated alginate

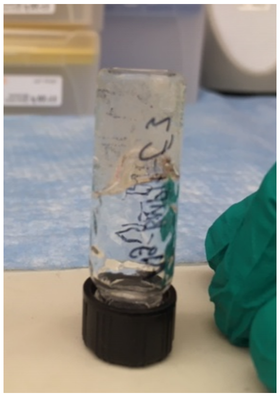

**Figure S9**: CC crosslinked alginate gels are self-supporting when inverted

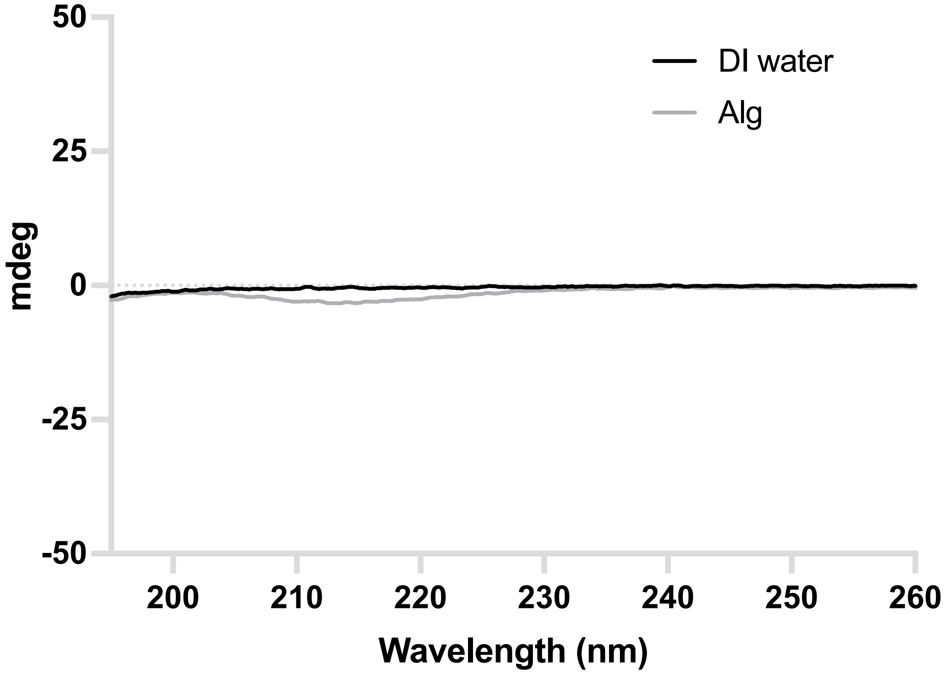


**Figure S10** Background spectra of DI $H_2O$ and 1% alginate in PBS

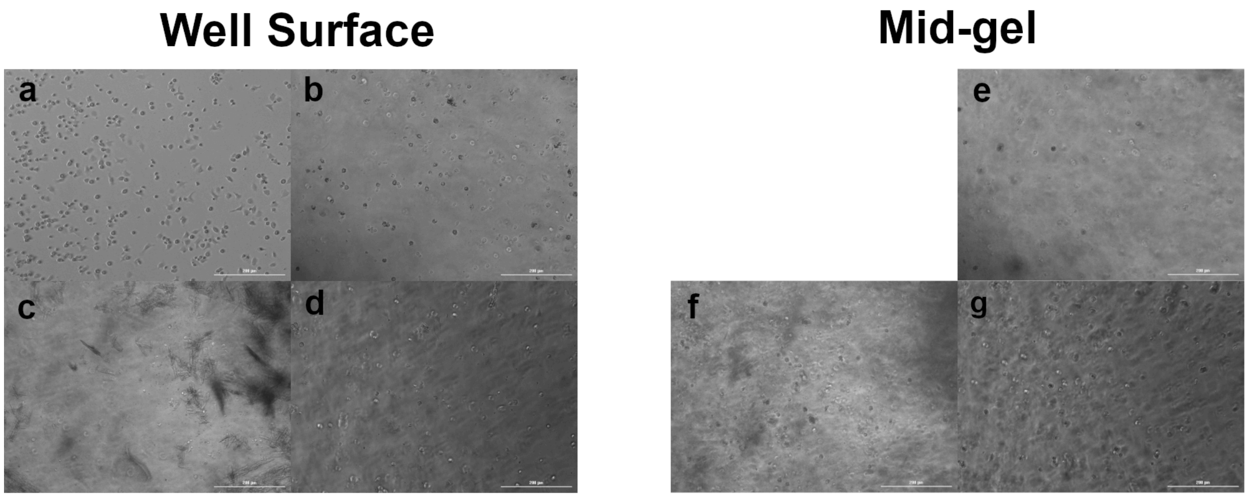


**Figure S11** (a-d) Primary MΦ were cultured for 24 hours either in tissue culture plastic (a) in $CaCl_2$ crosslinked gels (b) in $CaSO_4$ crosslinked gels (c) or in CC crosslinked gels (d) and images were taken at the surface of the well. (e-g) Primary MΦ were cultured for 24 hours in 3D

either in $CaCl_2$ (e) $CaSO_4$ (f) or CC (g) crosslinked gels and imaged within the bulk of the gel at 760 (e), 560 (f) or 420 (g) μm above the well surface.

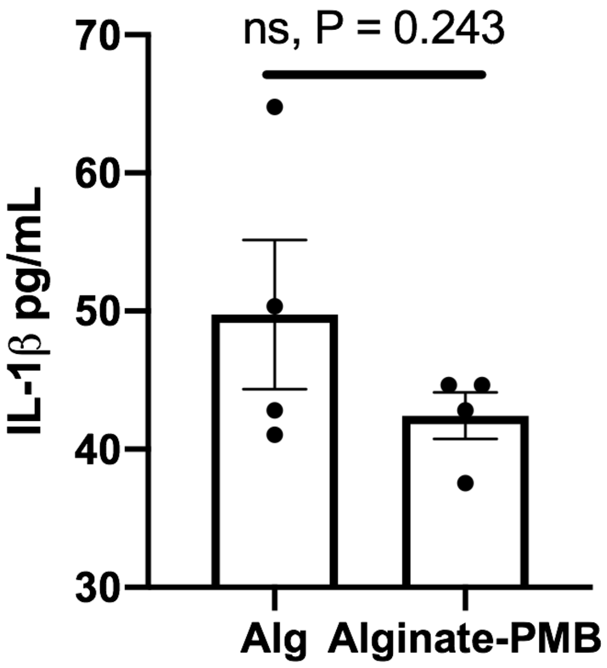


**Figure S12**. MΦ cultured in 2D exposed to 0.5% w/v alginate in the presence or absence of 10 ug/mL Polymixin-B and exposed to LPS at 500 ng/mL. No significant difference was observed in IL1-β expression by ELISA compared by students t-test.

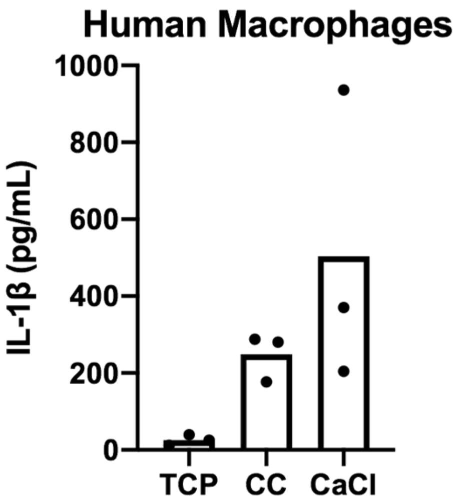


**Figure S13** Human PBMC derived macrophage IL-1β secretion in response to culture in CC or $Ca^{2+}$ crosslinked alginate culture.

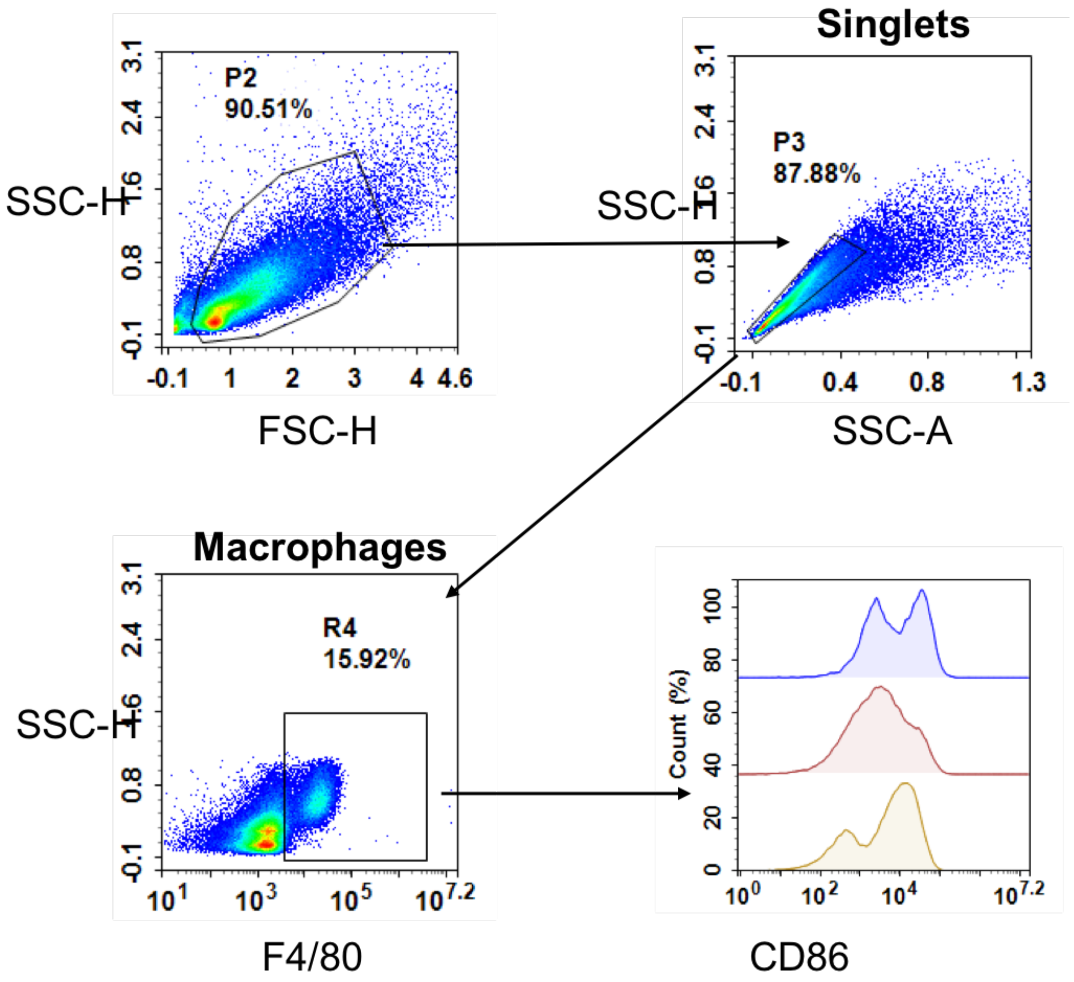


**Figure S14**. Gating strategy of macrophages exposed to $Ca^{2+}$ and CC crosslinked alginate with LPS stimulation *in vitro*, CD86 gating used as example.

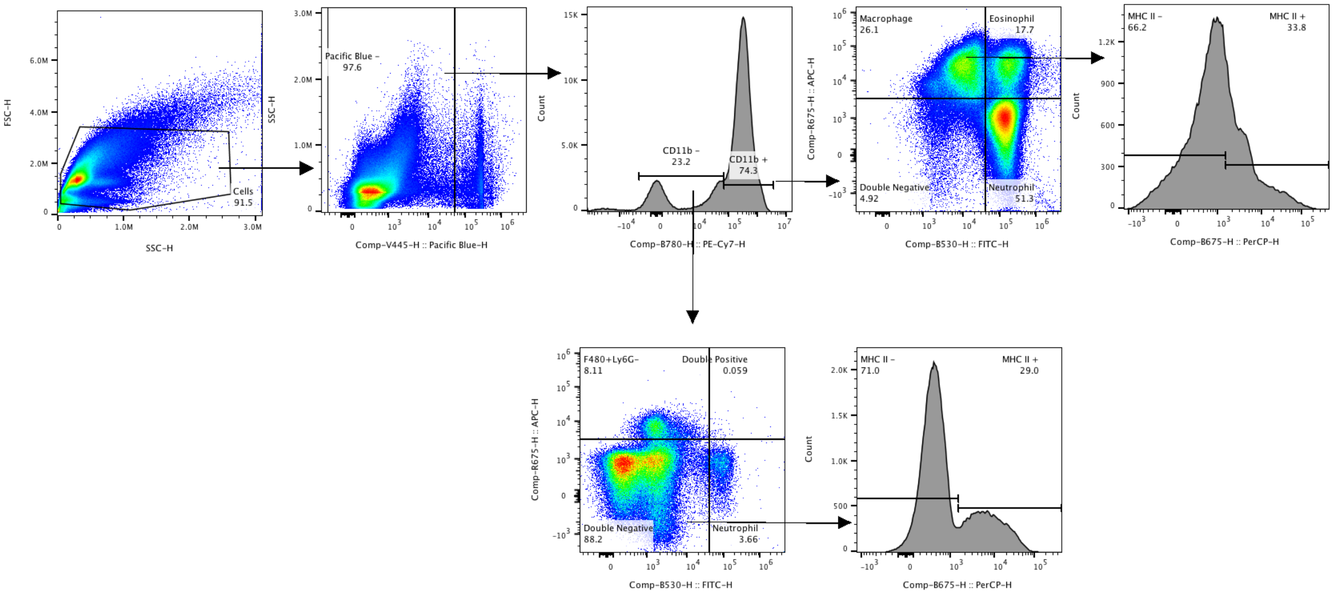


**Figure S15**. Gating strategy of peritoneal cells isolated from mice injected with either $Ca^{2+}$ or CC crosslinked alginate without additional LPS stimulation *in vivo*.

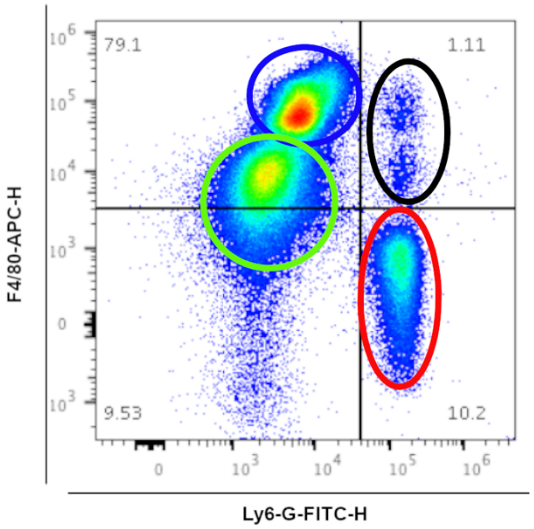


**Figure S16**. Four primary populations were observed in $CD11b^+$ cells. (Red) $CD11b^+$, $F4/80^-$, $Ly6\text{-}G^+$ cells were identified as $CD11b^+$ neutrophils. (Black) $CD11b^+$, $F4/80^+$, $Ly6\text{-}G^+$ cells were identified as eosinophils. (Green) $CD11b^+$, $F4/80^{int}$, $Ly6\text{-}G^-$ cells were identified as small peritoneal macrophages. (Blue) $CD11b^+$, $F4/80^{hi}$, $Ly6\text{-}G^-$ cells were identified as large peritoneal macrophages.

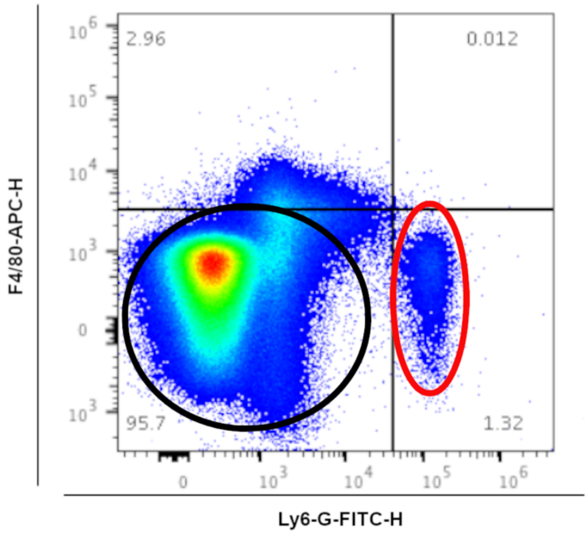


**Figure S17.** Two primary populations of were observed in CD11b$^-$ cells. (Red) CD11b$^-$, F4/80$^-$, Ly6-G$^+$ cells were identified as CD11b$^-$ neutrophils. (Black) CD11b$^-$, F4/80$^-$, Ly6-G$^-$ cells were identified as a heterogenous leukocyte population.
